# Oncogenic mutations of the EGF-Receptor ectodomain reveal an unexpected mechanism for ligand-independent activation

**DOI:** 10.1101/009068

**Authors:** Laura Orellana, Adam Hospital, Modesto Orozco

## Abstract

Epidermal Growth Factor Receptor (EGFR) signaling is supposed to be triggered by a dramatic ligand-induced conformational change of the extracellular domain from a closed, self-inhibited tethered monomer, to an open untethered state, which exposes a loop required for dimerization and activation. The ectodomain also untethers spontaneously, but the molecular mechanism is still unknown. Experiments with the mAb806 antibody have suggested the existence of a third untethered state, still uncharacterized, appearing transiently during the tethered to untethered transition, and which exposes a cryptic epitope hidden in the known structures. Here, molecular dynamics simulations of two ectodomain mutants (R84K and G39R) targeting a hinge at domain I-II interface highlight the possibility of such additional intermediate conformer. The new conformation is untethered but surprisingly compact, and originates from a large rotation of domain I that exposes the mAb806 epitope, in a similar way as the EGFRvIII deletion does. The present findings not only point to different molecular processes for ligand-dependent and spontaneous untethering, but also suggest that domain I-II mutation clusters and the EGFRvIII deletion may share a common structural mechanism, based on the removal of an steric hindrance from domain I that restricts dimerization.

## HIGHLIGHTS

> Mutations at the domain I-II interface favor spontaneous untethering

> The mutant untethered intermediate has a compact, not extended, configuration

> A backwards rotation of domain I exposes the mAb806 cryptic epitope

> Domain I rotation is sterically equivalent to the EGFRvIII deletion

## INTRODUCTION

Epidermal Growth Factor Receptor (EGFR) is the founder of the family of transmembrane *Tyrosine Kinase Receptors* (TKRs) that control cell growth and differentiation (Holbro and Hynes, 2004; Hubbard and Miller, 2007). In humans the HER (Human-EGF-Receptor) system includes four members: EGFR (HER1, ErbB1), ErbB2 (HER2/Neu), ErbB3 (HER3) and ErbB4 (HER4), which are often mutated in cancer (Riese et al., 2007; Roskoski, 2004; Rudloff and Samuels, 2010). All ErbB receptors have a ligand-binding extracellular region or ectodomain coupled to a single transmembrane tail and a cytoplasmic kinase. The ectodomain is composed of two tandem repeats of a ligand-binding subdomain (I / III) followed by a smaller cysteine-rich subdomain (II / IV). In the inactive states of the ectodomains for EGFR/HER1 (sEGFR) (Ferguson et al., 2003; Li et al., 2005; Ramamurthy et al., 2012), HER3 (Cho and Leahy, 2002) and HER4 (Liu et al., 2012a), a self-inhibitory *tether* in domain IV keeps hidden a *dimerization arm* in domain II, which is central for dimer formation and stabilization (see for example (Chung et al., 2010)). Crystal structures (Garrett et al., 2008; Ogiso et al., 2002) suggest that ligand binding favors a large conformational change, from the tethered, closed inactive monomer to an untethered, open back-to-back “*proud*” or “*heart-shaped*” dimer that is thought to represent the active state (*Fig. 1A* and *B*, respectively) (Lemmon, 2010; Schlessinger, 2002). In this process, the ligand binding subdomains rotate dramatically to simultaneously contact EGF, while the *tether* releases the *dimerization arm* to form key receptor-receptor interactions. Despite the huge effort focused on its structural characterization, the molecular mechanism for activation at the extracellular level is not fully understood, and the question whether the ligand triggers the experimentally observed transition to the open untethered state (Dawson et al., 2007; Mi et al., 2011) or there is a preexisting conformational equilibrium which is then biased by binding is still controversial. Recent simulations of HER4 (Du et al., 2012) support an induced-fit model where ligand binding drives a conformational change from the closed to an extended, open-like conformation with a free dimerization arm. There are however several observations that do not fit in a classical ligand-driven dimerization scheme. Experiments early shown that the tether only contributes 1–2 kcal/mol to stabilize the closed tethered state (Burgess et al., 2003; Ferguson et al., 2003); and thus, spontaneous untethering should be expected in a small population of receptors. It is also known that EGF contributes additively to dimer stabilization (Chung et al., 2010; Lemmon et al., 1997), and recently, it has been demonstrated that the biologically active species at the low EGF levels *in vivo* is probably a singly-ligated, not doubly-ligated dimer (Liu et al., 2012b), which requires ligand-independent monomer untethering. For these reasons, it is currently assumed that EGFR spontaneously samples *extended* untethered conformations which are then stabilized by the ligand (Bessman and Lemmon, 2012; Ferguson, 2009). However, in spite of data in favor of the preexisting tethered-untethered equilibrium (see for example (Chung et al., 2010; Kozer et al., 2011)), we cannot ignore that a spontaneous transition of a closed sEGFR monomer towards an extended form has still not been directly detected either in SAXS experiments (Dawson et al., 2007) or in microsecond-long simulations (Arkhipov et al., 2013).

**Figure 1.**
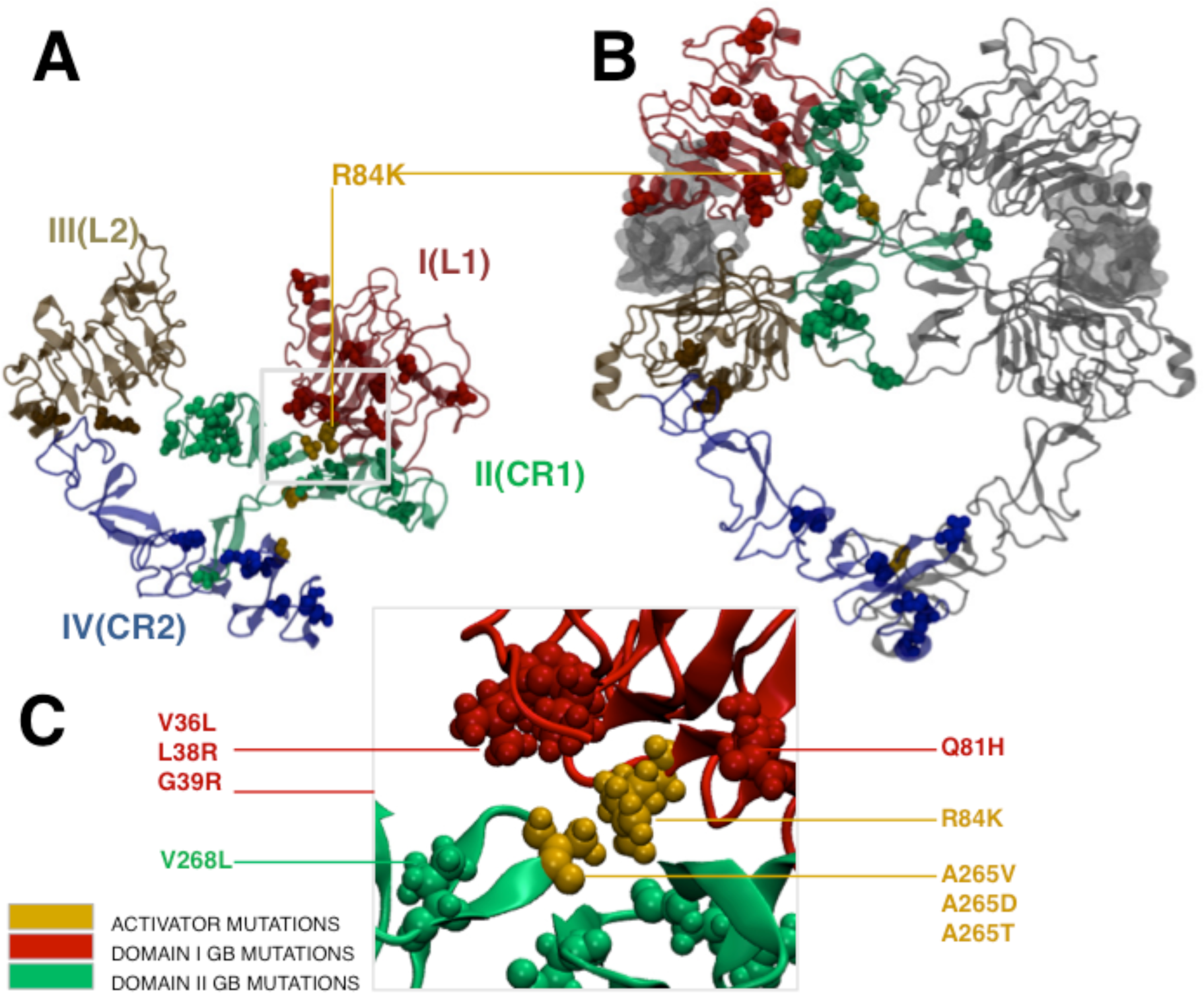
EGFR extracellular domains showing point mutations. (A) sEGFR inactive closed monomer, (B) active open dimer, (C) detail of domain I-II interface where most *glioblastoma* mutations are clustered. Known mutations are shown as balls, highlighting in yellow those known to increase sEGFR activity. Note the evident clustering of mutations at the interface between domains I (red) and II (green), and domains II and IV (blue), as well as the position of R84K mutation away from domain I-III ligand binding cleft (A) as well as domain II dimerization interface (B).

The analysis of available data on EGFR conformational landscape suggests that the open-close conformational transition might be more complex than anticipated. For example, a Cys-bonded module in domain II, inaccessible in the open and close structures, is known to be recognized by antibodies mAb806 and mAb175. Work by Scott and colleagues has demonstrated that these antibodies bind an intermediate form of the receptor after it untethers, but before forming the bound dimer (Johns et al., 2004). The mAb806 antibody was originally raised against the major glioma mutation, EGFRvIII, an N-terminal truncation of the entire domain I (Mishima et al., 2001). Since mAb806 epitope appears blocked by domain I in the close and open structures, but it is detected upon EGFR overexpression (Jungbluth et al., 2003) or under glycosylation (Johns et al., 2005), it became evident that a third untethered conformation must exist. The structure of this untethered transition state has eluded full characterization so far, and only the “cryptic” epitope module (C287-C302) has been crystallized in complex with the antibodies (Garrett et al., 2009).

An important but poorly explored source of information on the conformational equilibrium of sEGFR comes from glioma point mutations mapping the extracellular domain, not as common as the EGFRvIII deletion but also able to trigger ligand-independent activation (Huang et al., 2009; Lee et al., 2006). It is widely assumed that these *gain-of-function* mutations favor either the transition to the extended state and/or ligand binding (Rudloff and Samuels, 2010). Arguing in favor of a self-inhibitory role, a number of mutations target positions around the domain II-IV tether (57X-cluster in *Table S1*)), but interestingly, others concentrate at interdomain surfaces not immediately related with known functional regions, such as the domain I-II interface (3X- and 8X-clusters in *Table S1*, see *Fig. 1C*). These conserved positions, close in sequence and 3D-space, appear repeatedly mutated across different tumors, suggesting that they might affect the ectodomain dynamics.

Trying to cast light on the activating mechanism of I-II interface mutations, we performed a comprehensive study of the dynamics of Wild Type (WT) and mutated ectodomain using an Elastic Network Model (ENM) and Molecular Dynamics (MD) simulations. To get a preliminary view on the intrinsic flexibility of the tethered state, we first performed ENM calculations (Orellana et al., 2010). ENMs is a sequence-unspecific method that efficiently samples the major rigid body motions of a given protein architecture (Alexandrov et al., 2005; Bahar et al., 2010a, 2010b; Tama and Sanejouand, 2001). Analysis of ENM-predicted movements for the ectodomain suggested the domain I-II interface as a key hinge region for domain I oscillations. Based on these results we selected two different mutations (R84K and G39R) targeting the potential I-II hinge site for MD simulations. Both mutants showed spontaneous untethering of the ectodomain, but surprisingly, not to an extended, but to a highly compact conformation predicted by ENM. The new configuration is characterized by a backwards rotation of domain I-II that simultaneously exposes the dimerization arm and the mAb806-epitope in a similar way as the EGFRvIII deletion does, removing domain I from the putative dimerization interface. These results reveal an unexpected ligand-independent untethering process, and suggest that EGFRvIII and I-II point mutations may share a common structural mechanism, providing a molecular explanation for their similar biological profile.

## RESULTS

### Domain I-II interface acts as a hinge for domain I oscillations according to ENMs.

The structural transition between closed tethered (*1NQL* (Ferguson et al., 2003)) and open untethered (*3NJP* (Lu et al., 2010)) crystal states of sEGFR is dramatic (rMSD = 25.14 Å), and involves a series of rigid domain movements, with minimal local changes in all domains except domain II, which bends as a spinal backbone to allow dimerization. ENM analysis (see *Experimental Procedures*) suggests that in the tethered structure, domains I-II and domain II-IV form two rigid blocks, with domain I-II joint showing a greater flexibility. The ENM *normal modes* describe the possible concerted movements of the ligand-binding domains as these two blocks rotate or shift along an orthogonal plane dividing the structure (see *Fig. 2A*). The two softest modes (with a similar eigenvalue) trace back-and-forth oscillations of domain I against domain II as the I-II and III-IV planes twist (1^st^ mode), as well as domain I-III closure as they approach and bend the structure in a scissoring motion (2^nd^ mode); all these motions are accompanied by domain II bending or extension. Although most of the crystallographical *closed to open* change (75%) can be described by only ten modes, there is no mode dominance and the transition is distributed over several of them (1^st^, 3^rd^ and 4^th^, with overlaps ∼ 0.4), suggesting that it is not fully spontaneous. Residue fluctuations (see *Experimental Procedures*) show that the interacting dimerization arm (II) and the tether (IV) define the central rigid region (*Fig. 2B, green box*) that divides the structure into two blocks, and clearly point to the interdomain I-II region as a hinge-like surface (seen as a series of minima in domain I; *Fig. 2B*) for the oscillations of domain I over domain II (as seen in *Fig. 2A, modes 1-2*). An extreme number of glioma mutations (up to 15 point changes, see *Table S1*) exactly target the predicted I-II ENM-joint, which contains a series of small β-strands (Val36-Leu38, Glu60-Ala62, and Leu80-Arg84) forming hydrogen bonds and hydrophobic interactions with the opposing II surface. Overall, these results strongly suggest that particular I-II interface mutations can impact the native motions of the tethered state.

**Figure 2.**
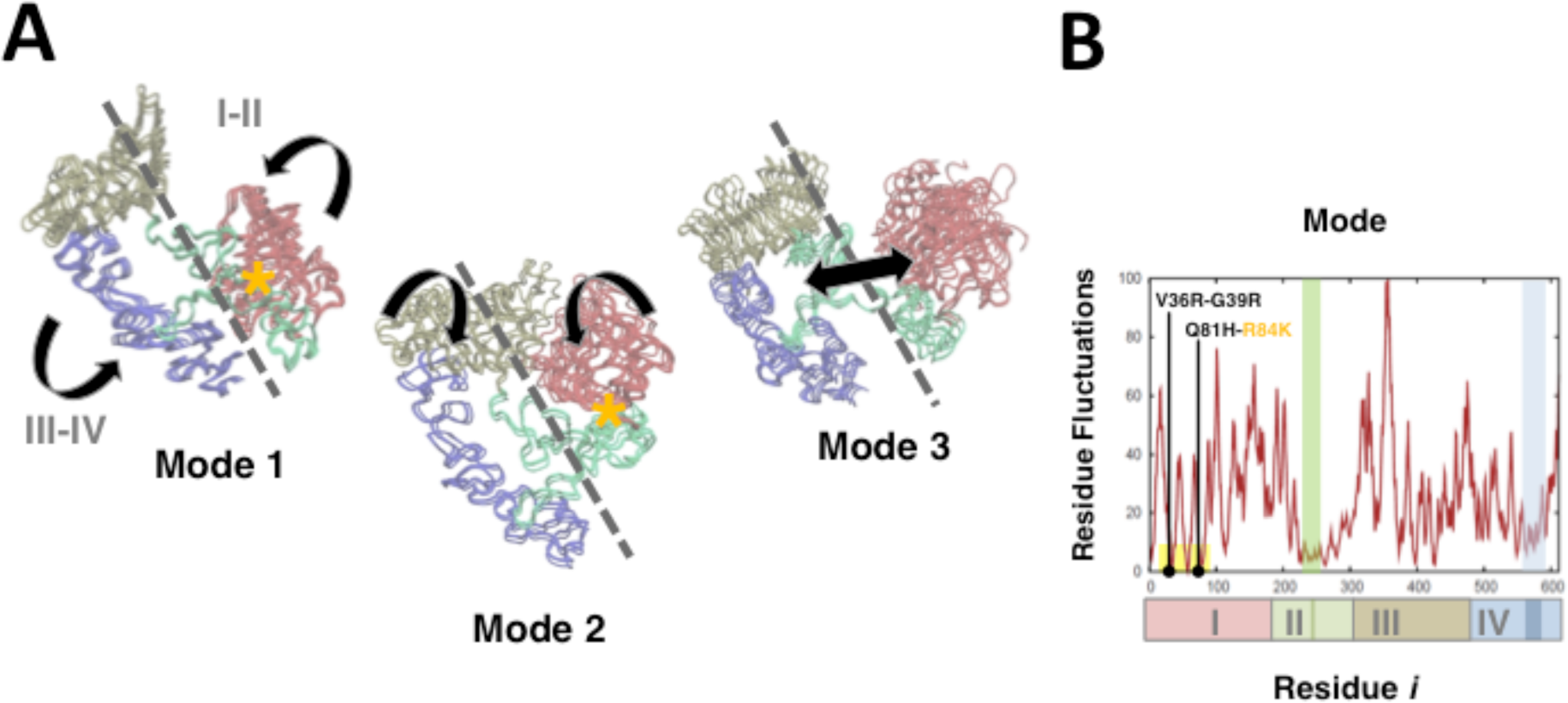
Global Flexibility of the ectodomain by Elastic Network Models (ENMs) (A) Structural ensembles along the first three normal modes of the closed sEGFR show domain I-III rotations. The directions of motion of the ligand binding domains (I and III) are indicated with curved arrows. Domains I-IV colored *as in Fig. 1*. (B) ENM Residue Fluctuations for the first 10 modes; rigid sites appear as minima and flexible regions as peaks. The dimerization arm/tether are marked with green/blue boxes; oncogenic mutation clusters in the I-II hinge surface (*highlighted in yellow*) shown as solid black circles.

### MD simulations show an extraordinary flexibility in closed sEGFR but no untethering.

In order to characterize the native WT ectodomain dynamics, we ran multiple unbiased runs of the tethered state obtaining 0.5µs metatrajectories (see *Experimental Procedures*). Trajectories of WT closed sEGFR show a pattern of motions in notable agreement with ENM-predictions (see *Table S2*), confirming that closed WT sEGFR samples spontaneously a wide range of conformations for the ligand-binding domains (*Fig. 3A-B*, note the wide rMSD distribution). This mobility is, as predicted, mainly due to oscillation motions of domain I over domain II (*Fig. 3A*, marked with arrows), where the interface acts as a hinge for domain I oscillations (*Fig. 3C*, highlighted in yellow). In some configurations, the domain I motions lead to dramatic increases in domain I-III distances, stretching domain II and loosening the tether-dimerization arm contacts (see *Fig. S1*, *green cluster 6*). All these movements are enhanced as a single trajectory is extended to the 0.5µs range (*Fig. 4A*), with the domain I-III cleft increasing up to 60Å (*Fig. S2A,* arrow in the 3^rd^ column), domain II stretching to near 180°, and the tether breaking transiently (see *Fig. S2A*, 2^nd^ and 3^rd^ columns arrow/star). After stretching, domain II relaxes and bends again, finally triggering a collapse of the ligand-binding domains at the end of the simulation (*Fig. 4A,* last frame) that greatly corresponds to the 2^nd^ ENM mode (*Fig. 2A)*. This remarkable stretching/bending mechanics of domain II, and the coupled oscillations of domain I, are mediated by the plasticity of the Cys-loop connecting the blocks formed by domains I-II with domains III-IV, as noted by Du and colleagues (Du et al., 2012) in HER4 simulations. However, in spite of the flexibility of the unbound tethered monomer, in the absence of ligand these spontaneous movements never approach an open-like or extended state, and the dimerization arm remains shielded by domain IV as other μs-long simulations have repeatedly reported (Arkhipov et al., 2013; Du et al., 2012). Interestingly, in all trajectories, the displacement of the tether results in crystallographically important tether residues (with the exception of HIS566) forming mostly intra-domain interactions, while residues in the mutated 57X-cluster (such as ALA573, GLY574 or GLU578) are readily engaged in a network of domain II-IV hydrogen bonds with mutated residues at the 23X- and 24X-clusters (such as TYR239 or TYR249) (see list in *Table S4*).

**Figure 3.**
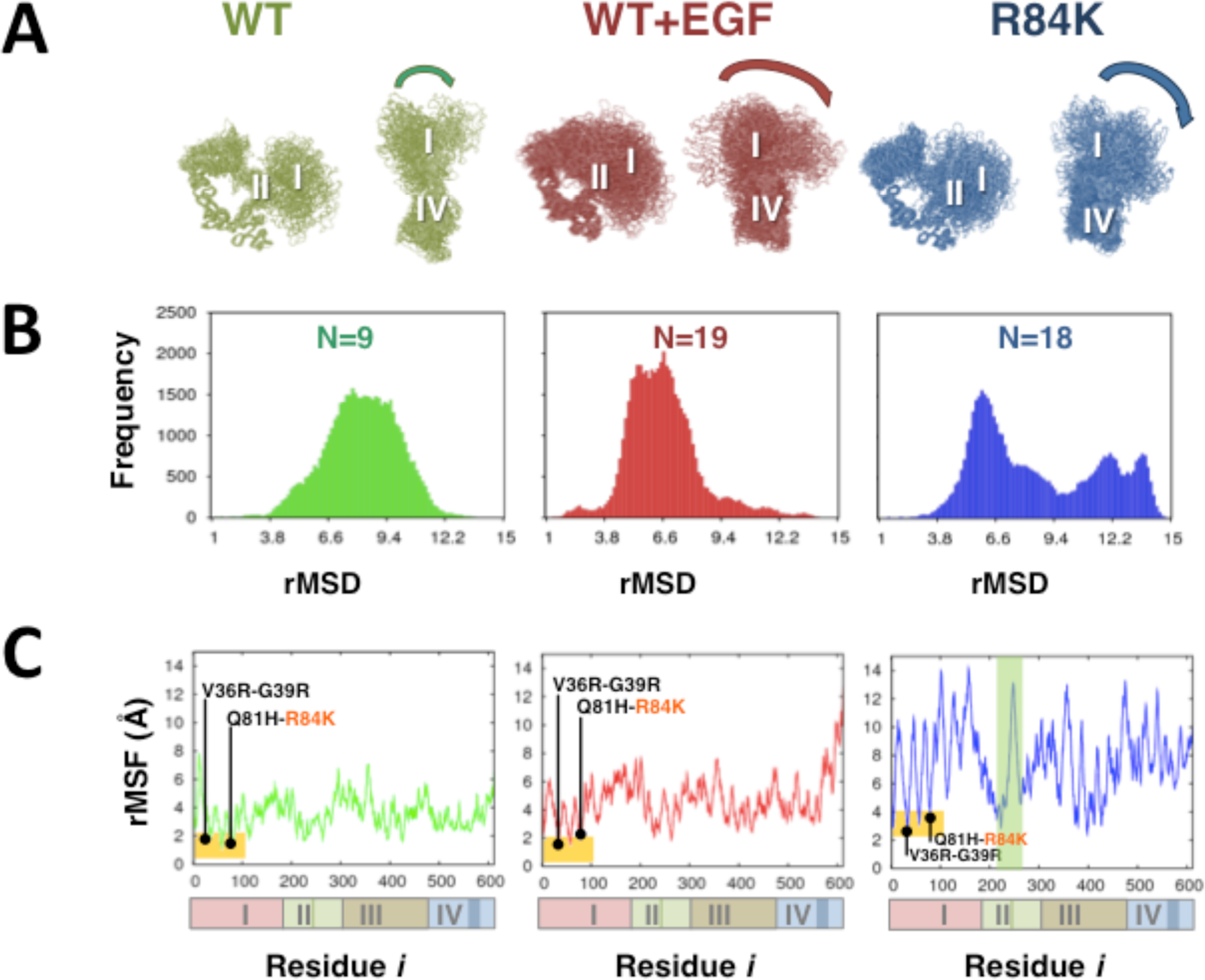
Analysis of 0.5µ s metatrajectories for the closed state. (A) Domain I oscillations in the metatrajectories, (B) Root mean square deviations (rMSD) distribution for MD snapshots with respect to the starting crystal structure (*1NQL*). Aligned clusters appear in *green* (Wild-Type, WT, unbound), *red* (WT+EGF, bound) and *blue* (R84K mutant, unbound); a view from the front (*left*) and the bottom (*right*) is shown, to highlight domain I oscillation movements (*curved arrows*). Compare the amplitude of domain I oscillations in the bound and R84K states with WT reduced motions. *N*= number of clusters. (C) Residue root Mean Square Fluctuations (rMSF). Domain I-II hinge points are highlighted in yellow, with the 3X- and 8X-mutation clusters shown as black dots. Note the increased motions of domain I in the R84K mutant.

**Figure 4.**
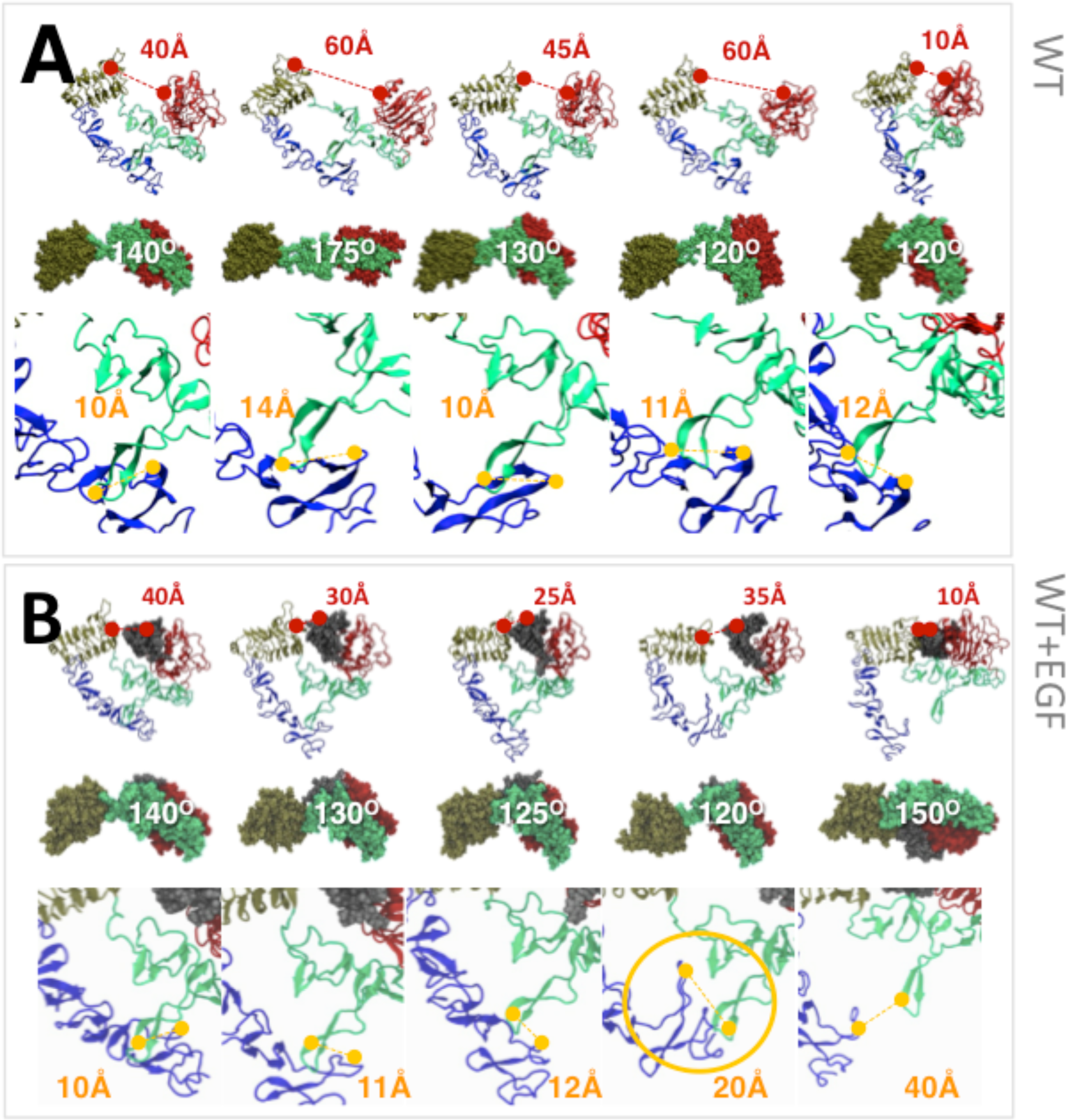
Submicrosecond Wild-type dynamics in the absence and presence of EGF. Representative structures obtained in very long MD trajectories for Wild-Type (WT) sEGFR alone (A), and in the presence of EGF (B). Domains I-IV colored as in *Fig. 1*. The starting state is the same for all simulations (*1^st^ frame*). For each box A-B: Front view of snapshots with domain I-III distances in red (*upper row*); a view from below to highlight domain II bending coupled to changing domain I-III distances (domain IV removed) with the bending angle in white (*middle row*); close-up to the tether region, II-IV distance in orange with irreversible untethering events highlighted with a circle (*lower row*). (A) WT sEGFR undergoes large domain II stretching motions (∼180°) coupled to domain I-III separation, followed by slow domain II relaxation. (B) Ligand-driven untethering follows domain I-III apposition and repeated domain II bending.

### Ligand binding drives the closed state to an extended untethered conformation.

In order to characterize ligand-driven untethering for HER1, we performed several simulations of the ectodomain with EGF bound. Following results by Du *et al.,* which showed that the ligand initially binds to domain I as a first step the activation process (Du et al., 2012), we simulated the binding pose found in the HER1 ectodomain crystal, observing a similar ligand-driven untethering process. Even in short MD runs, EGF binding to the WT closed state increases the alignment of the intrinsic dynamics along the *closed to open* transition (*Table S2*). The ligand remains bound to the binding site at domain I, shifting its oscillation motions over domain II to rotate and orthogonally approach domain III (*Fig. S1, red cluster 2*). As domains I-III get closer, domain II is forced to bend, contrary to the free stretching movements predominant in the unbound WT and R84K ectodomain (*see below*). After each bending event, relaxation of domain II separates again domains I-III. This alternate binding/unbinding of domain I-bound EGF to domain III, coupled to bending/relaxation of domain II, is followed by loosening of the tether, up to the point that transient untethering is detected even in short simulations (*Fig. S1 red cluster 13*).

A trajectory extended to 0.5 µs (*Fig. 4B*) clearly shows how the ligand triggers a periodic bending of domain II (see *Fig. S2B, 2^nd^ column grey arrows*) coupled to successively longer tether opening events. When irreversible untethering finally happens (*Fig. 4B*, 4^th^ frame, *yellow circle* and *also Fig. S2B*) domain III-IV quickly departs from domain I-II (see details in *Fig. 4B)*, leading to an extended-like conformation similar to the open untethered state of the crystal dimer (*Fig. 4B*, last frame). As observed for HER4 (Du et al., 2012), this *open-like* state still needs a rotation of domain III-IV to form the high-affinity binding site observed in the bound dimer, a step probably requiring dimerization (see *Discussion*).

### MD of the R84K mutant reveals a spontaneous transition to an untethered but closed state.

To test the hypothesis that mutations at the I-II interface may affect the dynamics of the tethered state as ENM suggested, we focused on the well-known activating glioma mutation R84K (Lee et al., 2006; Vivanco et al., 2012) for MD simulation. Despite the conservative nature of the Arg-> Lys change and its location away from the tether, closed R84K displays significant dynamical changes with respect to the WT ectodomain. In all four metatrajectory replicas a crucial hydrogen bond connecting Arg84 with the highly conserved and also mutated Cys227 is lost (*Table S3*), resulting in multiple subpopulations in the 0.5µs metatrajectory (*Fig. 3 top*). This augmented flexibility is due to a floppy I-II interface, which allows the mutant to display *enhanced* WT-like motions: there are large-scale oscillations of domain I over domain II (*Fig. 3A* and *Fig. 3C, yellow box)* leading to extreme domain I-III rearrangements and variations in the dimension of the binding cleft (note changing I-III orientations in *Fig. S1*), which are accompanied by stable untethering in ≈25% trajectories (2^nd^blue cluster in *Fig. S1*).

To analyze slow relaxation motions we extended again two of the R84K simulations to 0.5µs: while one rapidly reaches the same collapsed state as WT sEGFR (overlapping with the 2^nd^ ENM mode), the other displays fast but stable untethering. Analysis of this trajectory (*Fig. 5A*) confirms that spontaneous untethering is caused by maximization of the intrinsic closed WT motions, already predicted by WT MD and ENM simulations. The enlargement of domain I-III distance (near 60Å, *Fig. S2C,* 3^rd^ column arrow) stretches domain II (up to 180°, *Fig. S2C,* 3^rd^ column arrow, and *Fig. 5A,* 2^nd^ frame), but in the mutant, the domain I rotates more freely and tilts back up to contact domain IV breaking the tether (see *Fig. S2C star,* and *Fig. 5A*, yellow circle). This sequence of movements generates the complete exposure of the dimerization arm in an untethered conformation stable throughout the remaining 500ns simulation (*Fig. S2C*). A number of contacts between domains I and IV, such as a salt bridge between ASP147 and LYS569 (*Fig. S2D*) and further hydrogen bonds between ASP147 and THR569 side chains are formed upon untethering and stabilize this conformation. The spontaneously untethered unbound state in R84K trajectories has a compact, not extended, configuration (*Fig. 8A*), and the dimerization arm and the mAb806/175 epitope are both exposed. Although the diameter of the binding cleft is slightly narrower, EGF can be docked rendering stable bound complexes (see *Experimental Procedures*). We also tested the stability of the novel configuration for the WT receptor running several replicas with the original sequence; only in one case the structure re-tethers back to the closed state, confirming that spontaneous untethering is stable but also reversible.

**Figure 5.**
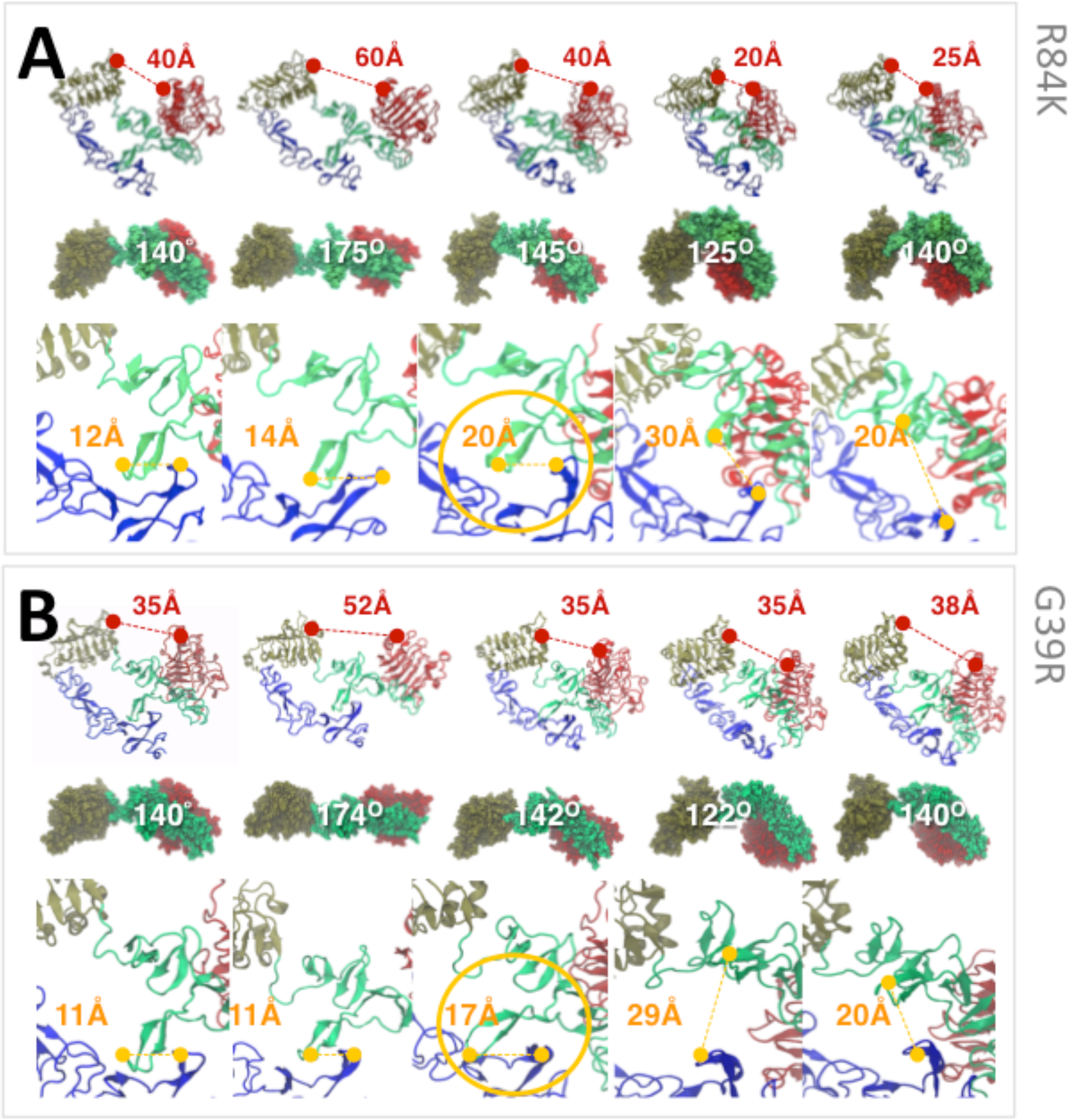
Submicrosecond dynamics for R84K and G39R mutations. Representative structures obtained in very long MD trajectories for the R84K (A) and G39R (B) sEGFR mutants. Spontaneous untethering in mutants follows domain I-III separation and domain II stretching as in WT trajectories, but here the extreme flexibility of the I-II interface traps domain I in a tilted-back configuration breaking the tether. As in the presence of EGF, the tether is broken when domain II returns to its native bending angle (∼140-150°, *1^st^ frames*).

### The energy landscape of the R84K simulations reveals two potential wells populated by oncogenic mutants.

In order to obtain a picture of the global flexibility of the unbound tethered state we performed PCA of the WT and R84K metatrajectories considered together (1μs), which cover a broad range of conformations (see *Experimental Procedures*). The two major Principal Components (PCs) of the combined meta-trajectory show excellent overlap with ENM modes (*Fig. 6A),* and clearly separate the crystal structures solved for the HER Family into its different members and conformational states (*Fig. 6B, black triangles*). Using PC1 and PC2 as reaction coordinates, we also estimated the free energy profile (*Fig. 7A-B*), clearly different in unbound WT and R84K. While simulations of the WT ectodomain remain in the same energy minimum as the closed structures solved (*Fig. 6B and Fig. 7, left)*, the R84K mutant samples more extensively the conformational space along both PC1 and PC2. In particular, R84K visits a region of the 1^st^ principal component out of the WT basin never captured crystallographically, visible as a large cluster corresponding to the untethered intermediate (*Fig. 6B right, cluster 2, in yellow)*. This cluster corresponds to a second local minimum (*Fig. 7 right),* which is connected to the native basin by a shallow well. The energy barrier between both basins, corresponding to untethering, is below 2kcal/mol, consistent with previous estimations (see *Discussion*). The R84K mutant also explores a wider region at the upper boundary of the native basin (*Fig. 7 right)*, corresponding to the domain I-III collapsed state along the 2^nd^ principal component (*Fig. 6B right, in green),* only sampled by the WT ectodomain in the 500ns simulation.

**Figure 6.**
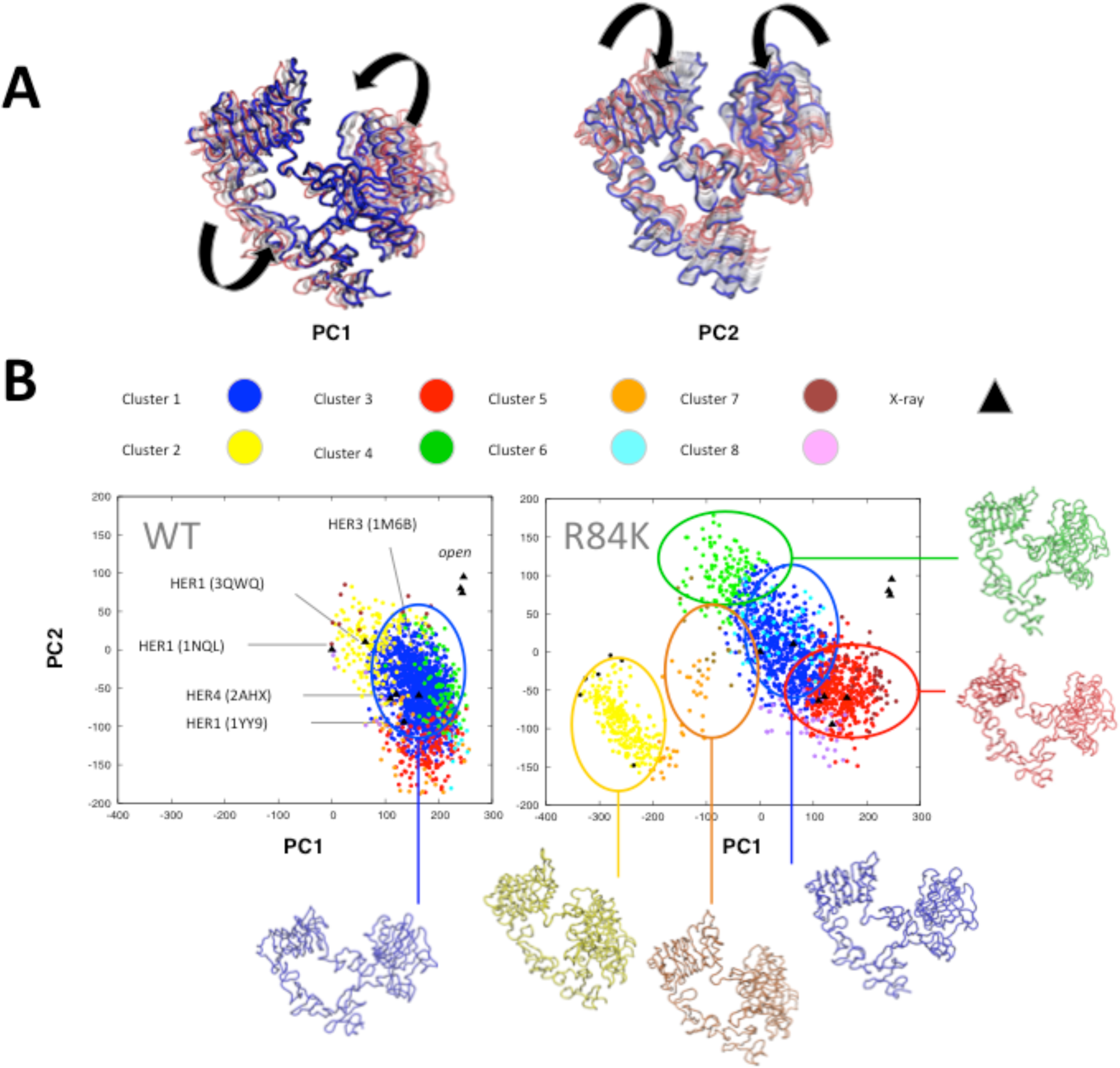
(A) First two Principal Components (PC) of the WT and R84K metatrajectories considered together. The directions of motion of the ligand binding domains (I and III) are indicated with curved arrows. (B) Projection of the major clusters onto PC1-PC2. Clusters are colored according to their relative populations for either WT or R84K metatrajectories. The intermediate state sampling PC1 corresponds to the second cluster in R84K (*in yellow*).

**Figure 7.**
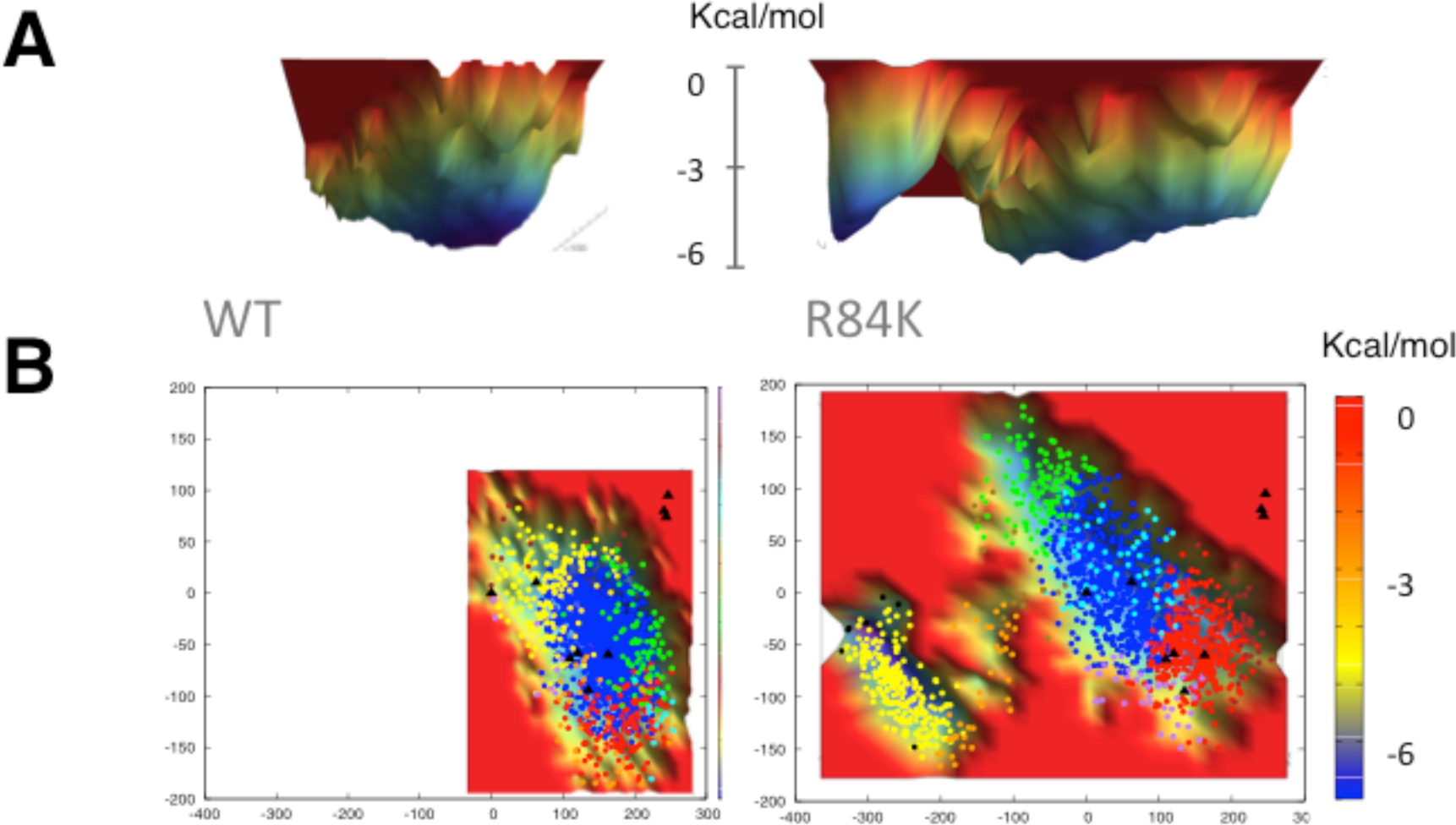
Free energy landscape of WT (left) and R84K (right) unbound ectodomains. (A) Energy surface (view from below). (B) Contour map showing the projection of the clusters in *Fig. 6*. Reaction coordinates are defined by the two Principal Components (PC1-PC2) from the conformational space sampled by MD metatrajectories considered altogether (1μs). Note how the mutant ectodomain explores a second local basin along the first Principal Component (PC1) corresponding to the untethered intermediate.

None of the trajectories approaches the open configurations represented by the HER2 monomer and the HER1 bound dimer, which are out of the main basin (right upper corners in *Fig. 6B*).

### Simulations of the nearby mutation G39R reveal a similar transition to a compact untethered state.

As a final test for the statistical significance and reproducibility of the observed transition, we ran independent MD simulations of the glioma mutation G39R, which is located in the close vicinity of R84K. The oncogenic activity of this mutation has not been studied *in vitro,* as is the case for R84K, but it is right in the center of a mutation cluster (V36R, L38R/P, G39R/W) not only for glioma but also lung and colon cancer. In the case of G39R, the mutation introduces a charge in a patch of hydrophobic interactions between domains I and II. Quite encouraging, we observed in one of the G39R replicas a virtually identical untethering transition early in the simulation (*Fig. 5B)*, and resulting in a mAb806 epitope-exposed compact untethered state (see *Fig. S3A*) stabilized by a similar network of I-IV interactions (*Fig. S3B*). This agreement between independent simulations of two different mutations suggest that ectodomain oncogenic mutations might share a common pathological mechanism, related to the modulation of the conformational landscape of sEGFR.

## DISCUSSION AND CONCLUSIONS

In comparison to the large number of computational studies devoted to the EGFR kinase domain, the dynamics of the extracellular domain and the effect of its mutations have received little attention to date. In spite of recent efforts to simulate the ectodomain (Arkhipov et al., 2013; Du et al., 2012), key aspects of the activation mechanism at the extracellular level are still poorly understood. It is has been widely assumed that, either triggered by ligand binding or spontaneously, untethering and receptor opening are parallel processes; in other words, that untethered configurations are extended. For that reason, activating mutations are generally supposed to spontaneously shift the ectodomain towards open untethered states, although there is no direct proof supporting that hypothesis. Here, we provide evidences that interdomain I-II mutations might shift instead the dynamics towards an untethered but *closed-like* state.

Our initial ENM analysis and WT MD simulations showed that while the essential deformation movements of unbound sEGFR align well with the open to close transition, spontaneous opening is unlikely in the absence of ligand. Such results are in agreement with previous simulations (Arkhipov et al., 2013; Du et al., 2012) and EM data (Mi et al., 2011), which show that the unbound ectodomain preferentially adopts compact conformations. The ENM-analysis also indicated that the natural motions for closed sEGFR are large-scale rotations of the ligand-binding domains as the two blocks formed by domain I-II and III-IV move in a concerted fashion. A rotational dynamics of the unbound ligand-binding domains is supported by data from solution spectroscopy (Kozer et al., 2011). Most interesting, ENM and PCA of the WT simulations pointed to the domain I-II interface, where a number of mutations concentrate, as a hinge surface for domain I oscillations.

The atomistic simulations showed that, while the unbound ligand-binding domains display significant rotational motions, the transition towards extended conformations is observed neither in WT nor in mutant R84K/G39R. However, R84K and the nearby I-II interface mutation G39R can spontaneously untether in the absence of ligand, acquiring a previously uncharacterized closed configuration, with domain I tilted backwards and downwards up to contact domain IV as in the 1^st^ ENM mode. Discrete domain I oscillations are detectable in crystal structures of the HER Family (note how they distribute along PC1 in *Fig. 6B*), but not to the extent and in the direction displayed by I-II interface mutants. Besides the feasibility of such motion according to crystal structures, several experimental evidences that cannot be explained with the two known conformational states support the existence and biological significance of this untethered intermediate.

*First*, the mentioned cancer-specific monoclonal antibodies (mAb806 and mAb175) which target a module in domain II inaccessible in both closed-tethered and open-untethered sEGFR (Gan et al., 2012; Johns et al., 2004; Jungbluth et al., 2003). Such cryptic epitope is known to be exposed only transiently during the conformational change from the tethered inactive to the untethered active state (Johns et al., 2004) but it is fully accessible in the mutant intermediate, making easy for docking and MD simulations to obtain stable sEGFR-antibody complexes (see *Fig. 8B*) which agree with crystallographic and mutagenesis data (*Fig. S5A-B*) (Chao et al., 2004; Garrett et al., 2009). Mutations targeting the Cys-loop connecting domains II-III have already been tested for mAb806 binding (Ymer et al., 2011), demonstrating that an increased flexibility at this region boosts the recognition of the cryptic epitope. This observation is also in agreement with our mutant simulations, where the flexibility of the II-III hinge clearly can facilitate the backwards rotation of domains I-II.

**Figure 8.**
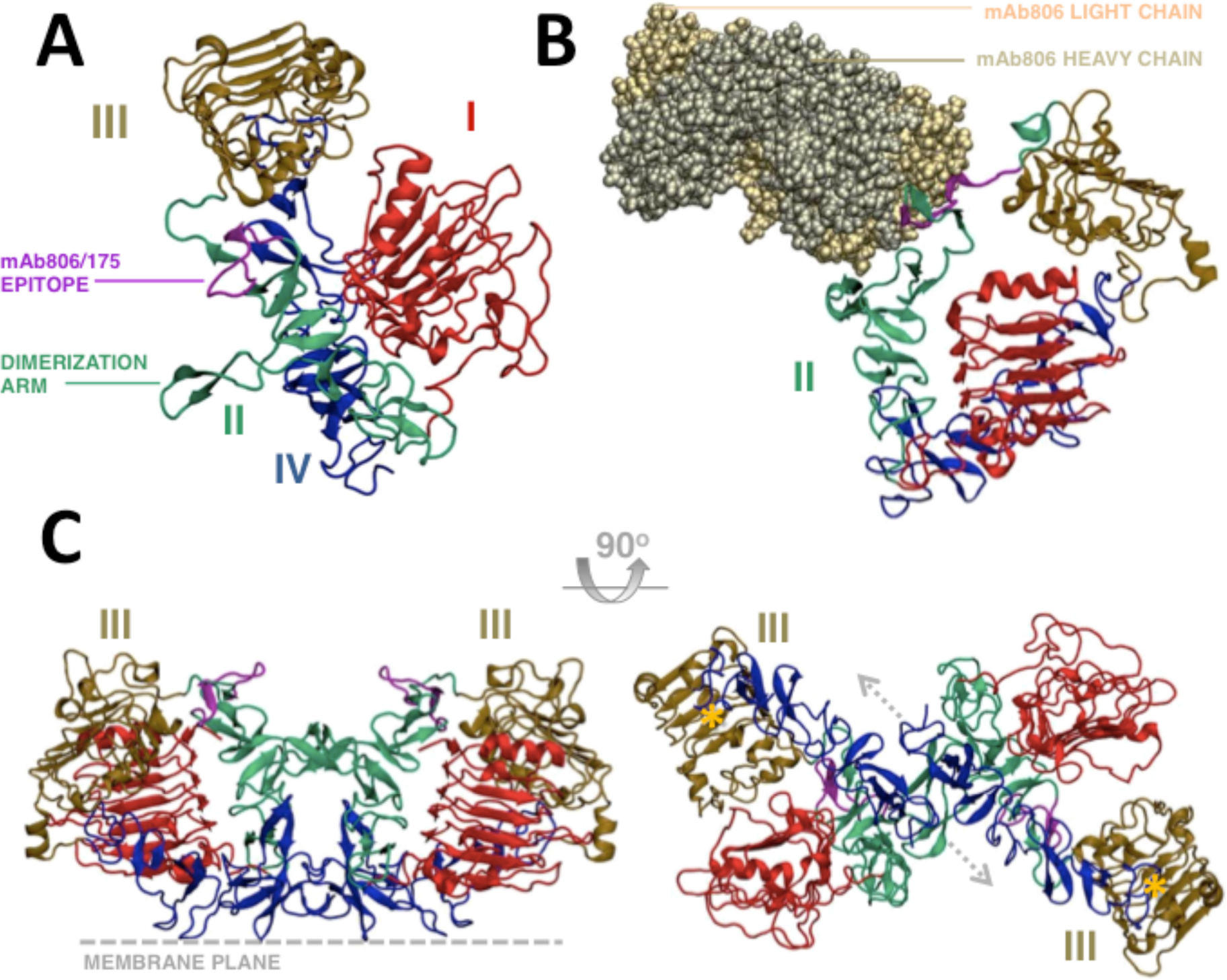
Ectodomain R84K untethered state. A) Domain disposition of sEGFR after the untethering transition. Note how the extreme backward rotation of domain I, up to interact with domain IV, allows full access to the mAb806 cryptic epitope (*highlighted in purple*) as well as the dimerization arm (*green*). B) Putative model of the complex of the antibody mAb806 bound to the R84K mutant intermediate state. This figure shows the intermediate bound to antibody (PDB: 35GV, chains show as yellow-ochre filled spheres) after MD relaxation. C) Hypothetical “Crouched” unbound dimer resulting from direct docking of two intermediate monomers: front (*left*) and bottom view (*right*); note the anti-parallel disposition of domain IV-IV (*dashed arrows*) and the large domain III-III distance *(right, marked with yellow stars),* and how the cryptic epitopes (*purple*) remain exposed. The plane of the membrane passes under the structure in the front view (*dashed line*) and parallel to the paper in the bottom view.

*Second*, the closed-like conformation of this novel untethered state could help to understand SAXS results (Dawson et al., 2007), which revealed subtle conformational changes (actually, a more compact conformation) upon tether weakening mutations, but not a transition towards the expected extended state. If untethering in the absence of ligand leads to closed-like configurations of sEGFR similar to the intermediate observed, instead of extended structures, minor changes should be expected in low-resolution experiments. Once again, this is in agreement with the above mentioned trend of the unbound ectodomain to retain compact configurations, even in the absence of the tether (Liu et al., 2012a).

*Third*, it is very suggestive that the backwards and downwards rotation of domain I-II in R84K/G39R intermediates removes the domain I steric blockage on the mAb806 epitope in a similar way as the EGFRvIII deletion does (*Fig. 9A-B*). We noticed that while domain I is displaced downwards in the intermediate state, in the extended conformations is instead displaced upwards (see *Fig. 9C*). This suggests that displacing domain I from the inactive state position could be important to allow the approach and proper antiparallel orientation of the transmembrane (TM) and juxtamembrane (JM) segments required for kinase dimerization (Endres et al., 2013; Jura et al., 2009; Red Brewer et al., 2009). Up to date there is no satisfying explanation for the occurrence of both EGFRvIII and point mutations in gliomas, or for their oncogenic mechanism at the structural level. Assuming that both mutation types are sterically equivalent regarding domain I hindrance at the dimerization interface would help to understand their similar oncogenic phenotype (Rojas and Bögler, 2011) and response to kinase inhibitors (Vivanco et al., 2012).

**Figure 9.**
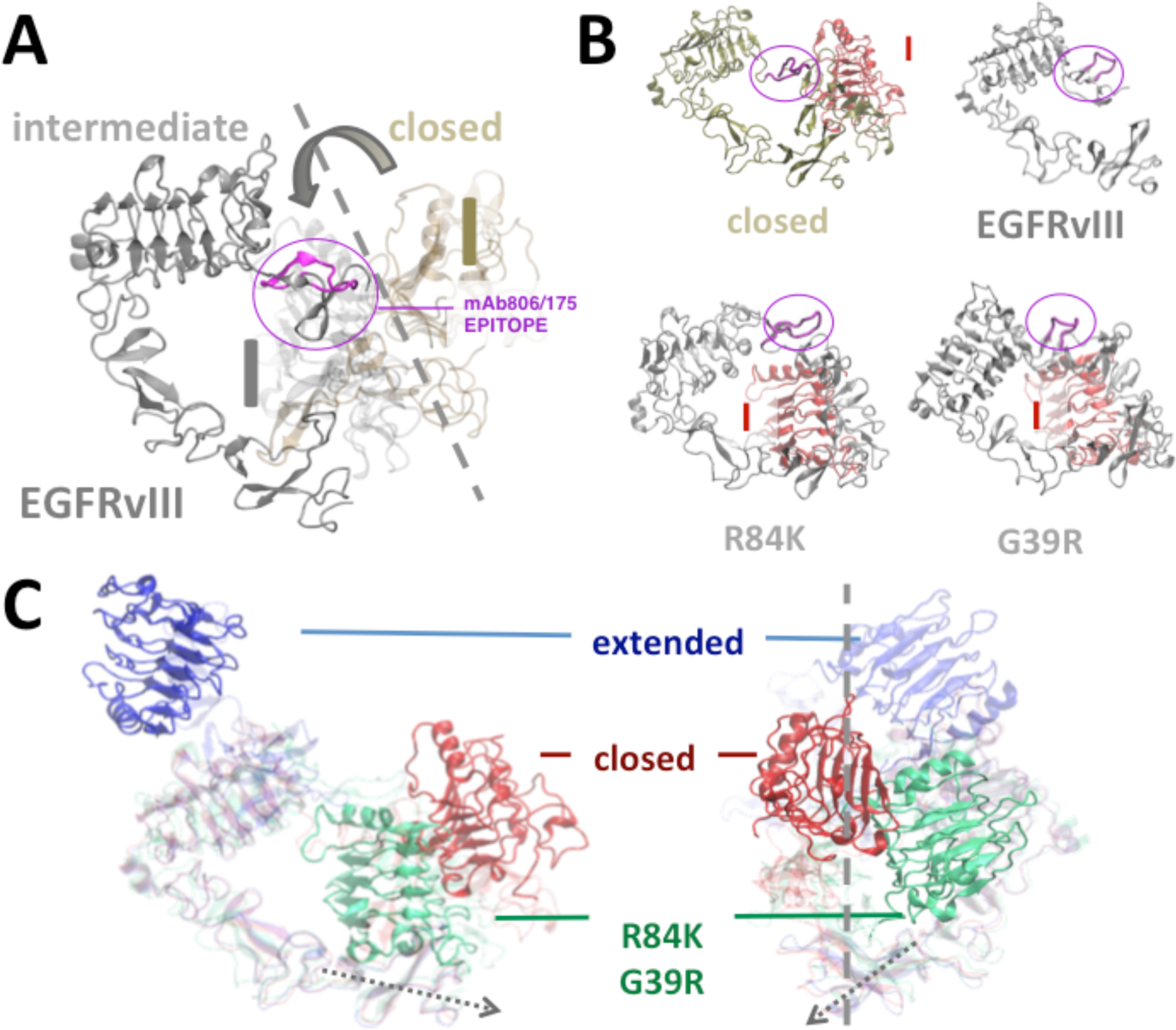
Structural similarity of R84K/G39R intermediates and EGFRvIII deletion. A) Comparison between domain I deletion in EGFRvIII (solid grey), domain I rotation in the intermediate (light grey) and domain I position in the closed state (light brown). The mAb806 epitope is shown in magenta. The deletion in EGFRvIII and rotation in the intermediate removes domain I from the closed position (highlighted with a dashed line). B) Structural equivalence between domain I deletion and back rotation in R84K/G39R intermediate. View from below showing how domain I (in red) sterically restricts the mAb806 epitope (magenta) recognition in the closed state, but not in EGFRvIII and I/II interface mutants. B) Position of domain I in the extended (blue), intermediate (green) and closed (red) configurations of the ectodomain. A R84K intermediate and the extended (*3NJP*) and closed (*1NQL*) conformers are shown aligned on the rigid domains III-IV in a front (left) and lateral (right) view. For clarity, only domain I is shown opaque for each configuration while the rest of the structure is colored transparent. Note how domain I is removed from the putative dimerization interface where the C-terminal tails meet the membrane by rotating up (in the extended state) or down (in the mutant intermediate).

*Finally*, a flexible sterical coupling between domain I orientation and the TM-JM segments could naturally account for the ectodomain dimerization promoted by kinase inhibitors. It has been shown that stabilization of the inactive configuration of the kinase by quinazoline inhibitors promotes the formation of unbound dimers that bind mAb806 (Gan et al., 2007) and are characterized by heterogeneous EM profiles clearly different from those of bound dimers, which are more similar to the X-ray “proud” configuration (Lu et al., 2012). Interestingly, the closed untethered state here reported seems perfectly designed to form a flat unbound dimer through the interaction of the dimerization arms (see *Fig. 8C*, details in *Experimental Procedures*) in excellent agreement with these data. The resulting dimeric structure can be described as “*crouched*” or “*rod-shaped*”, has an antiparallel disposition of both domains IV approaching their C-terminal ends (*Fig. 8C right*) and is still fully capable for the interaction with mAb806 (*Fig. 8C left*). Note that the distance between domains III-III in an intermediate dimer model (110-120Å, see *Fig. 8C right*) is much larger than in the X-ray “*proud*” configuration (70-80Å) but fits perfectly with that reported for kinase-induced dimers (118 ± 25 Å) (see (Lu et al., 2012)). Since the intermediate state is rather flexible, kinase-induced dimers, that appear as polymorphic cross-like structures when bound to Cetuximab, can be easily reproduced by R84K/G39R dimers (compare models for PD168393-cetuximab-EGFR in (Lu et al., 2010) with *Fig. S4A-B*).

Our simulations also point that WT sEGFR can also have a very small tendency to spontaneously untether towards closed-like states. Given the similarity of WT and R84K intrinsic motions, it is likely that the WT ectodomain also samples closed-untethered structures in longer timescales or under special conditions. A ligand-independent untethering process (see *Fig. 10*) running in parallel to ligand-driven untethering for WT HER1 could naturally explain the formation of both unbound dimers and singly ligated dimers (Liu et al., 2012b). In this scenario, free untethered monomers could act as *ready-to-dimerize* partners for the ligand-untethered ones, while the unbound pre-active dimers could also act as first target for ligand binding (as suggested by (Teramura et al., 2006)), forming in both cases singly-ligated dimers. In the presence of oncogenic mutations, the populations of untethered monomers and unbound dimers would increase dramatically, enhancing basal kinase activity and EGF-response as experiments indicate (Lee et al., 2006).

**Figure 10.**
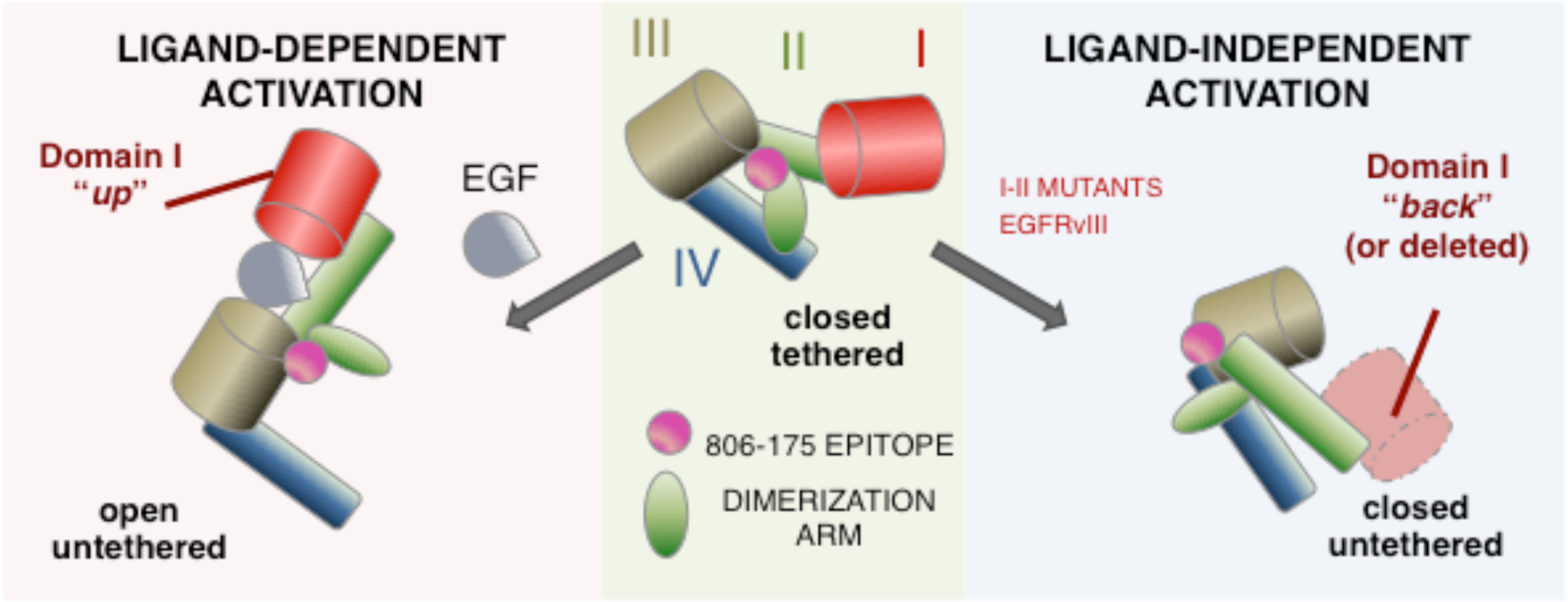
Proposed pathways for sEGFR activation. The scheme shows domain I arrangement and exposure of the dimerization arm (*green*) and the cryptic epitope 806/175 (*magenta*). To allow for dimer formation the domain II dimerization interface must be free from domain I sterical hindrance. In ligand-dependent activation domain I is moved to an “upwards” position by the ligand (*left*), while in the ligand-independent process (*right*) moves spontaneously backwards (or is deleted, in EGFRvIII) exposing the mAb806 epitope. In both cases the dimerization arm becomes untethered allowing further dimer stabilization.

Overall, these results suggest that the mechanism for mutation-driven activation might not be only the tether exposure itself, but also the acquisition of a orientation for domain I compatible with productive kinase dimerization. Under such scenario, the frequent convergence of point mutations and the domain I deletion (EGFRvIII) – which either acquire the intermediate configuration or mimic it-could be understood as a response to an evolutionary pressure favoring dimerization and activation of particular signaling pathways. The complex picture that emerges goes beyond the induced-fit/conformational selection dilemma, suggesting that mechanistically different processes may converge or be required to create functional dimers: a spontaneous one, towards a closed-like untethered state with domain I “*down*”, or a ligand-triggered one, towards an open-like untethered state with domain I “*up*”, allowing in both cases the proper interaction through domain II dimerization arms and the antiparallel TM/JM segments. Whether the observed spontaneous transition is only found in gliomas expressing domain I-II interface mutations or it represents a more general mechanism in other mutations and/or the WT receptor certainly requires further investigation, but certainly suggests the existence of multiple routes for EGFR activation, which could be sensitive to differential ligand concentrations or post-translational modifications affecting interdomain flexibility.

## EXPERIMENTAL PROCEDURES

### Structural data and naming scheme for extracellular domains of HER1.

We follow the naming scheme for sEGFR, distinguishing four CATH (Orengo et al., 1997) subdomains: ligand-binding domain I (residues 1-190), cysteine-rich domain II (191-309), ligand-binding domain III (310-481) and cysteine-rich domain IV (482-614). Our references are the 612-long fragments Glu3-Thr614 from closed (PDB: *1NQL*) and open (PDB: *3NJP, chain A*) ectodomains.

### Elastic Network-Normal Mode Analysis.

In the *Elastic Network Model* (ENM), the potential energy (*E*) is defined by a network of interactions with nodes at the C^α^ atom pairs *i, j* coupled by elastic springs, *K_ij_* (Atilgan et al., 2001; Tirion, 1996):

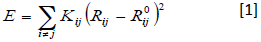

where *R_ij_* and *R_ij_^0^* are the instantaneous and equilibrium pair distances. In the normal mode analysis (NMA) approach (Case, 1999) the Hessian matrix of second derivatives is diagonalized, yielding 3N-6 eigenvectors or normal modes. The lowest-frequency modes represent the collective functional motions (Tama and Sanejouand, 2001). Here we use the nearest-neighbors MD-derived ED-ENM potential for distance-dependent springs (Orellana et al., 2010).

### MD Simulations.

Given the heterogeneity of glycosylation patterns and the poorly-defined orientation of the ectodomain in the membrane, we decided to simplify the system and neglect both the glycosyl moieties and the lipid bilayer as in previous simulation studies (Arkhipov et al., 2013; Du et al., 2012), in order to get clearer insight into the intrinsic large-scale flexibility of the extracellular domain. Thus we simulated WT (unbound and bound) and mutant R84K ectodomains starting from the closed *1NQL* configuration. Proteins were titrated, neutralized, hydrated, minimized, heated and equilibrated using standard protocols (Hospital et al., 2012). Initially, four replica trajectories for the three conditions studied (WT, WT+EGF, R84K) were collected during 50ns using the AMBER99SB-ildn force-field with GROMACS (Hess et al., 2008; Lindorff-Larsen et al., 2010). Two of the trajectories were extended to 0,2μs (closed); combined with 50ns replicas we obtained 500ns (closed) metatrajectories. Finally, two of closed state 200ns WT/R84K simulations were extended to 0,5μs. The control simulations with the ligand bound to domain I as in the *1NQL* crystal were set as in (Du et al., 2012) to check if the reported HER4 ligand-induced untethering also occurs in HER1.

Further simulations of the intermediate were performed to validate the feasibility of the untethered configuration for WT sEGFR, and its ability to bind ligand, the mAb806/175 antibodies or another untethered monomer: *a) Intermediate sEGFR WT:* we took a representative untethered configuration reached by the R84K mutant and reverted it to the WT sequence; *b) Intermediate sEGFR ligand binding in R84K and WT:* EGF was docked to domain I of WT and R84K intermediate states of sEGFR using as binding pose that one from the *1NQL* structure; *c) Intermediates EGFR-Antibody complexes:* mAb806 and maAb175 complexes with the sEGFR epitope were taken from PDB coordinates 3G5V and 3G5Y respectively and used to guide docking of the exposed epitope of our putative R84K intermediate; *d) Intermediate sEGFR dimers*: we used the *3NJP* dimer to guide docking of the dimerization arms exposed in two intermediate structures, obtaining an unbound “crouched” homodimer without steric clashes. The obtained structures/complexes (a-d) were subjected to MD relaxation (five 50ns replicas) to check their stability. To prove the significance of the observed transition, we performed further MD simulations of the mutation G39R in the close state. Total simulated time is 4.6μs, details in *Table S6*.

### Analysis of MD trajectories.

The noise arising from short-range vibrations was filtered by *Essential Dynamics*, ED (Amadei et al., 1993) to obtain Principal Components (PCs) representing collective movements. We used *in house* software, AMBER and GROMACS utilities to compute PCA, rMSD, rMSF, cluster analysis and the free-energy landscape; hydrogen-bonds and salt-bridges were analyzed with VMD plugins. To monitor structural changes, we used Cα-Cα definitions as in ENM:

*a) Domain II curvature:* was defined by the angle between Cα positions at the basis of the dimerization arm and the two extremes of domain II (residues 190-260-309)*; b) Domain I-III distance:* in the unbound state, was monitored by the Cα-Cα distance for the salt bridge pair LYS13-ASP364, which appears in both R84K and WT sEGFR simulations, and in the bound state, was monitored by the Cα-Cα distance for the EGF-sEGFR salt bridge pair LYS48-ASP323, which approaches domains I-III surfaces through bound EGF; *c) Domain II-IV tether distance* was monitored by the Cα-Cα distance between the hydrogen bonded pair TYR251-GLU578; d) *Domain I-IV distance in intermediate states:* was measured by the Cα-Cα distance for the salt bridge pair ASP147-LYS569, which appears in R84K and G39R simulations after untethering. Other angle and bond definitions yield similar results to track domain rearrangements.

### Metrics to analyze protein motions.

We used different metrics to study the protein motions from PCA and ENM modes:

*a) Similarity of protein motions* from either ENM or MD (described by a set of modes, i.e. eigenvectors, *v*_*k*_, and eigenvalues, λ_*k*_) between them and with experimental motions was computed as:

- Overlap of computed motions (from ENM or ED) with observed X-ray conformational change:

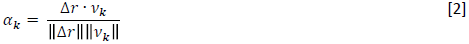

where *Δr = (R_2_ − R_1_*) / ‖*R_2_ − R_1_*‖ is the unitary transition vector between the two sets of coordinates, *R_1_* and *R_2_*, describing the observed states of the protein (closed and open) and *v*_*k*_ is the *k^th^* essential deformation mode. Generalization of *Eq.[2]* for the *m*-important deformation modes (i.e. explaining a given threshold of structural variance) yields a similarity index ranging from 0 (no similarity) to 1 (perfect similarity) (Tama and Sanejouand, 2001; Yang et al., 2009):

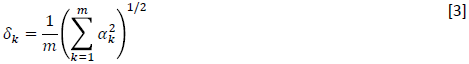

-Overlap between the motions computed from ENM and MD: Hess’s metric (Hess, 2000) was used to estimate the similarity of ENM and MD first *m*-important modes:

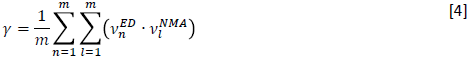

where the indexes *n* and *l* stand for the orders of the eigenvectors (ν, ranked according to their variance i.e. amplitude of motion and decreasing energy) and as in *Eq.[2]*, *m* is the number of essential modes considered.

b) *Residue flexibility:* the average thermal fluctuations for each residue *i* were computed from the ENM modes as in (Atilgan et al., 2001; Van Wynsberghe and Cui, 2006).

## ACKNOWLEDGMENTS

We thank the financial support of the MINECO (BIO2012-32868, and Consolider E-Science), the European Research Council (ERC-Advanced Grant), the Scalalife Project, and the Generalitat de Catalunya (SGR_2014). The authors thank Erik Lindahl for critical reading of the manuscript and helpful suggestions. Calculations have been carried out at Barcelona Supercomputing Center.

---

**Figure S1.**
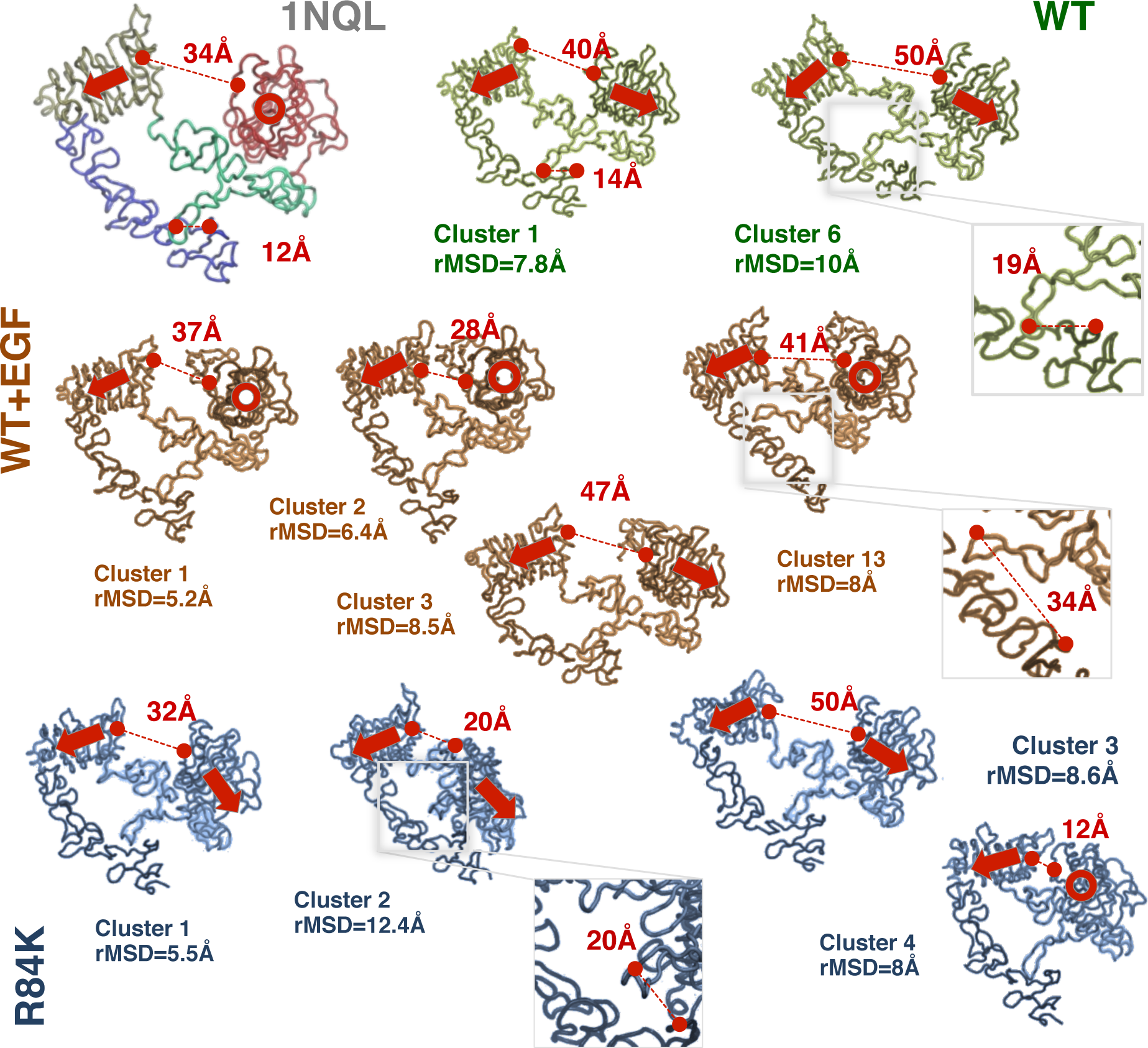
Major clusters and spontaneous untethering in closed state 0.5 µ s metatrajectories. *WT=Wild type sEGFR (green structures); WT+EGF=sEGFR with EGF (ochre structures); R84K mutant =sEGFR (blue structures).* The starting structure (1NQL) is shown as reference with domains I-IV red-green-ochre-blue. Interdomain Cα-Cα distances between domains I-III and domains II-IV appear in red (*dashed lines*); see *Methods* for definitions. The orientation of domains I and III are highlighted with arrows (N to C-tal); a circle indicates an arrow perpendicular to the page). Mutant and bound states are very polymorphic, with twice the number of clusters of WT sEGFR. Ligand-independent untethering is observed after domain I-III extension in WT state (*cluster 6*, detail) as well as in R84K (*cluster 3*), but in the last proceeds fast to an intermediate domain I-tilted state (*cluster 2*, detail) or to a collapsed I-III state (*cluster 4*). On the contrary, ligand binding approaches and orientates domain I orthogonal to domain III instead of parallel (*clusters 1-2*).

**Figure S2.**
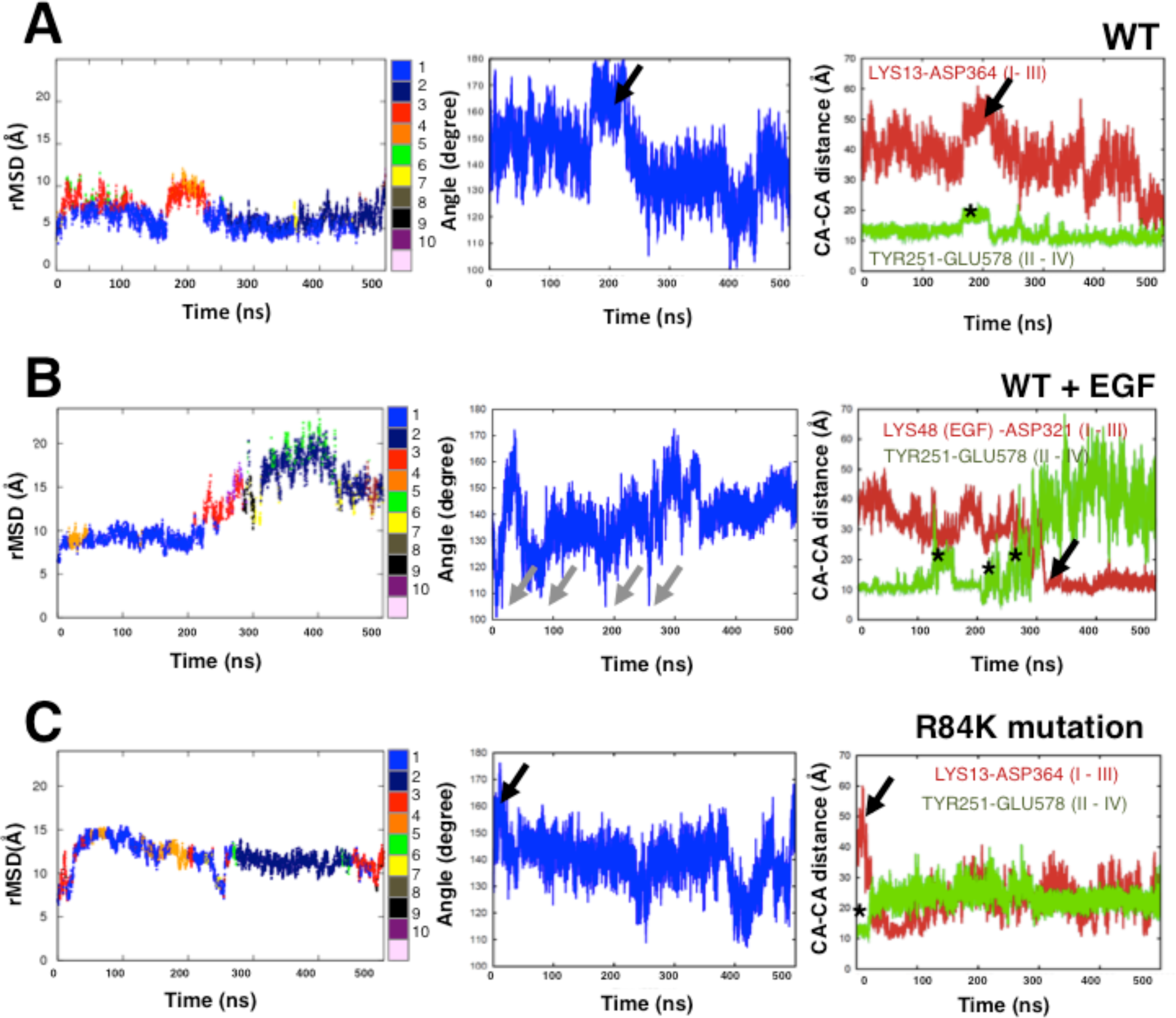
Extended 0.5 µ s trajectories for WT unbound sEGFR (A), WT bound sEGFR (B) and R84K unbound sEGFR (C) From the first to the third column, geometrical descriptors are: rMSD versus initial state colored according to major clusters (1-10), domain II angle bending, and domain I-III and II-IV distances (see *Methods* for definitions). Untethering events are marked with stars in column 3 (*green plot*). Note how in WT (A) and R84K (C) domain II stretching peaks (black arrows in column 2) are associated with domain I-III separation (arrows in column 3, *red*) and breaking of the tether. Ligand binding (see B) forces domain II bending rather than extension (grey arrows in column 2), which is followed by increasing tether breaking events after a certain relaxation time (*column 3*).

**Figure S3.**
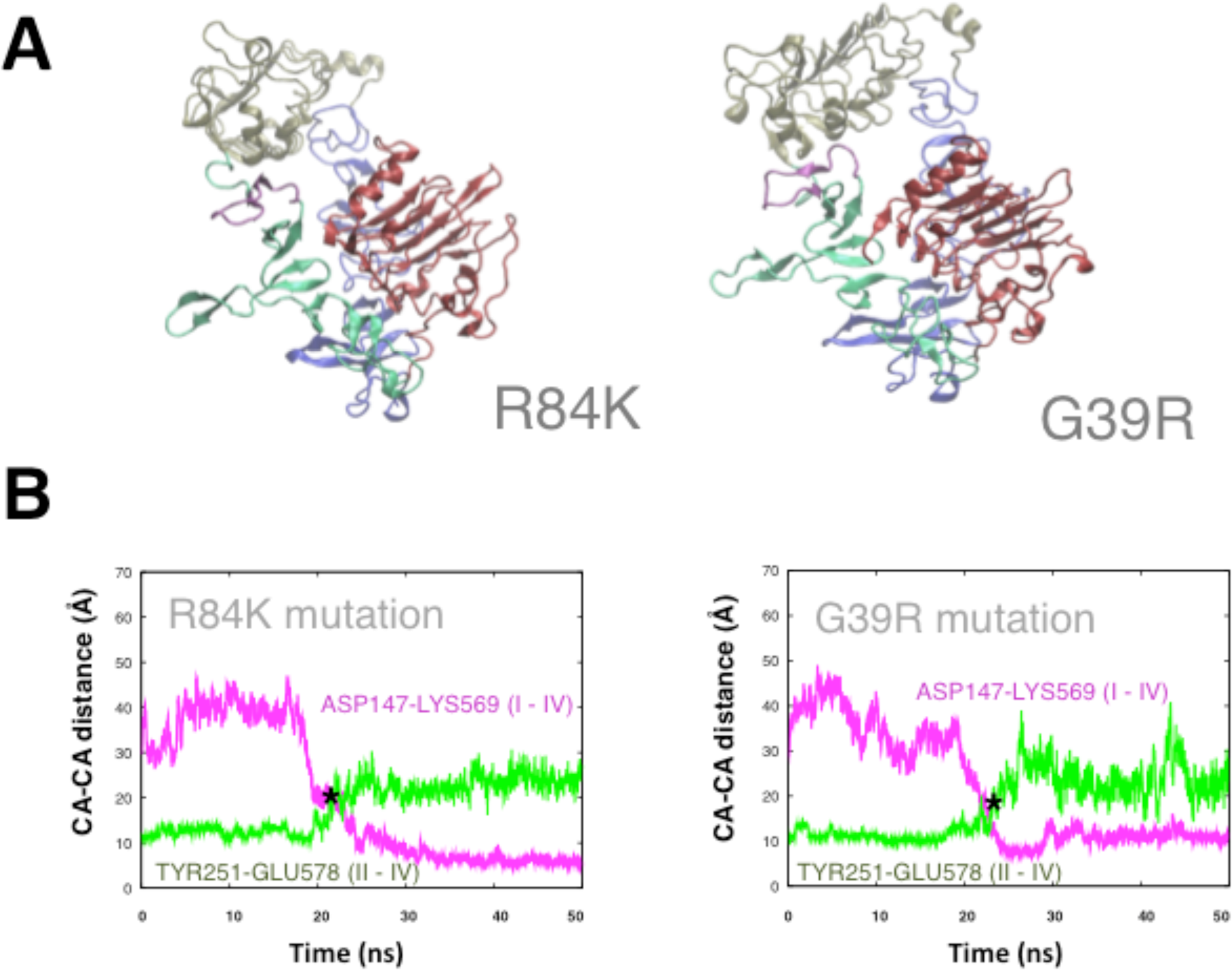
Spontaneous untethering in R84K and G39R leads to a similar compact conformation. A) Comparison between representative snapshots of the intermediate state for R84K and G39R sEGFR mutants. **B) Comparison of fast untethering events in two R84K and G39R trajectories.** The time evolution of the distances between domains II-IV (tether, in green) and I-III (in purple) is shown.

**Figure S4.**
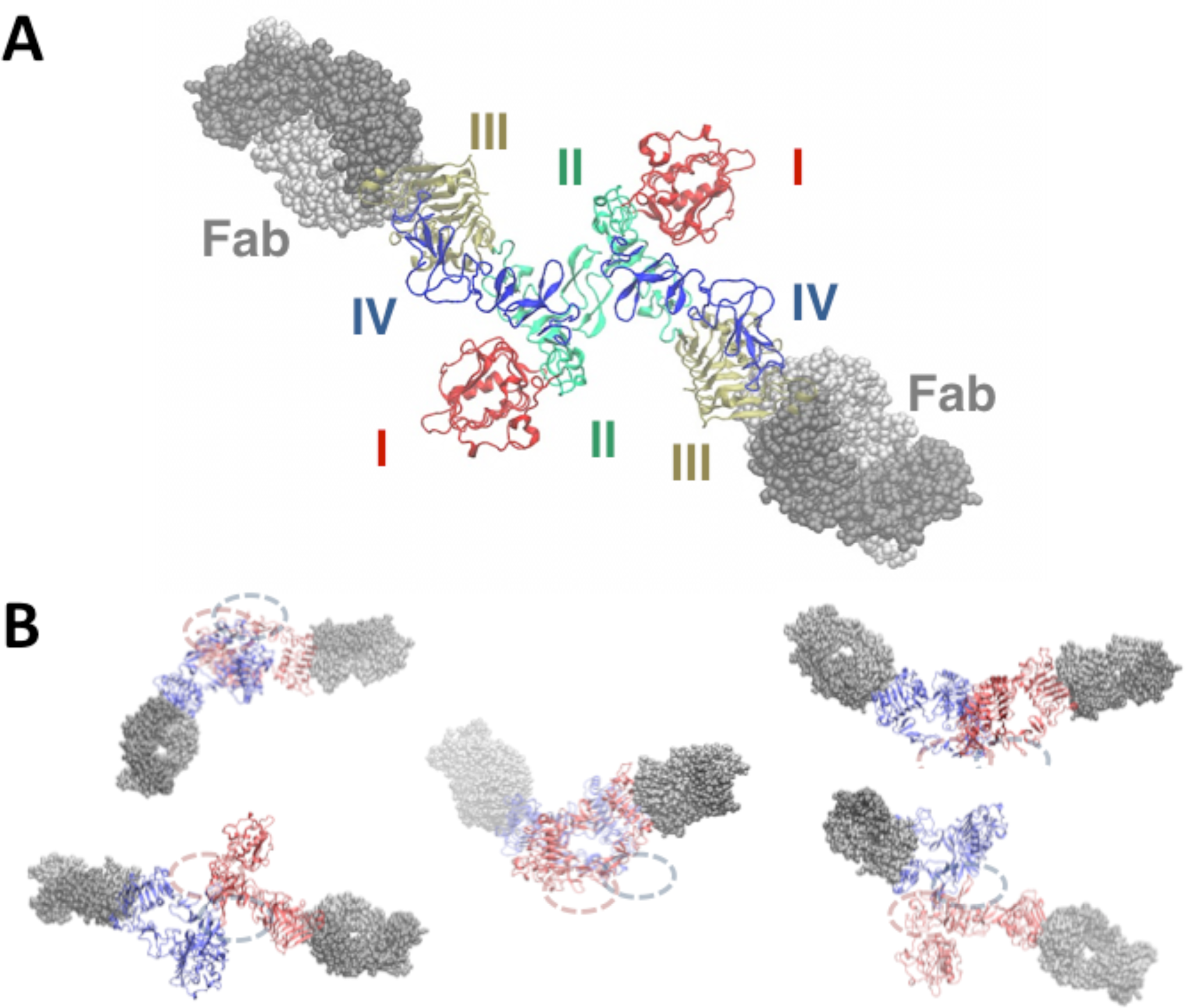
Intermediate dimers bound to Cetuximab. A) Intermediate dimer model generated from a G39R intermediate structure showing the orientation for each monomer of domain III bound to Cetuximab Fab. **B)** Different views of G39R and R84K dimers bound to Cetuximab (in grey) showing the approximate positions of the kinase domains for each monomer (pink and blue ellipses). Cetuximab Fab is docked to domain III as in the reference structure *1YY9*.

**Figure S5.**
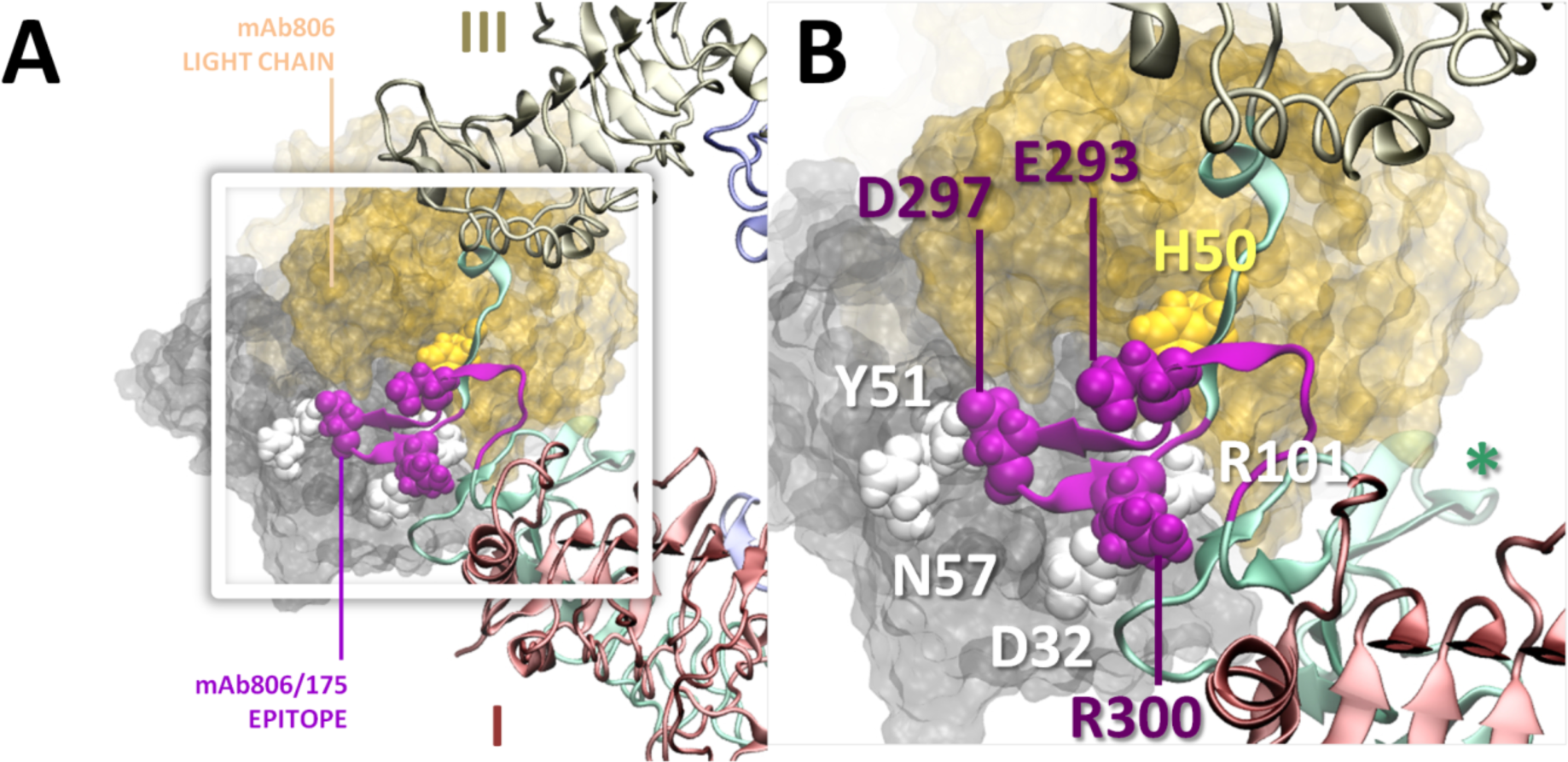
Two levels of zooming showing the interaction between the mAb806 antibody and the R84K exposed epitope: (A) Overall view showing domain disposition and (B) Detail of the antigen-binding site. This figure shows one of the possible models in which the epitope can be inserted into the antigen-binding site by docking to the crystallographic complex (PDB: 35GV), followed by minimization. The ectodomain backbone is shown as cartoon, with domains colored pink (I), green (II) and tan (III), and the epitope loop in purple; the antibody is shown as a surface with colored light (yellow) and heavy (gray) chains. Known interacting residues (E293-H50/R101, D297-Y51/N57, and R300-D32) are highlighted as white (heavy chain), yellow (light chain) or violet (EGFR epitope) spheres. Note the dimerization arm (star), away from the binding region in (B).

**Table S1.**
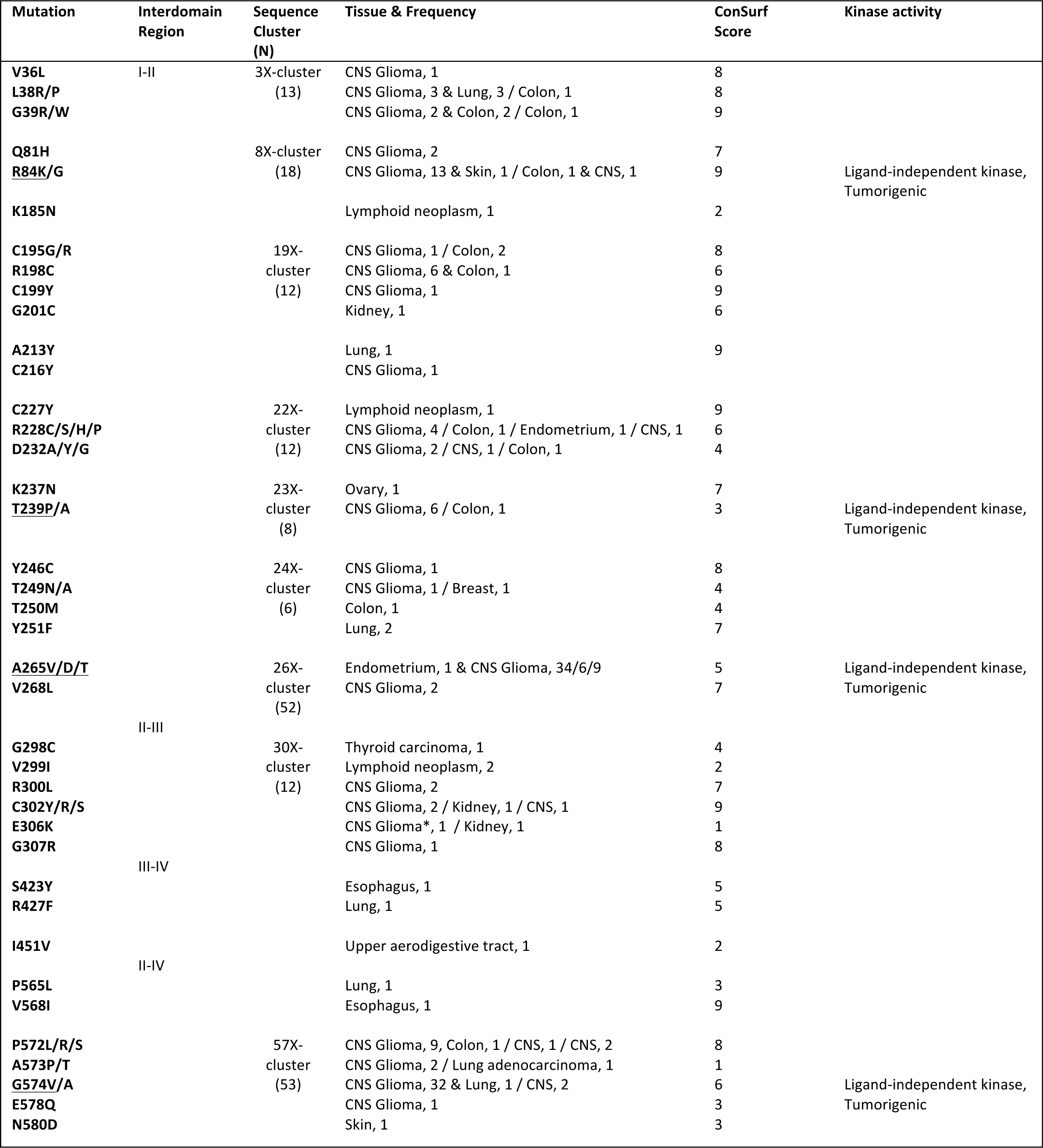
Summary of isolated and clustered missense mutation in the EGFR ectodomain targeting interdomain regions, as registered in the COSMIC database. Residues are considered cluster neighbors up to three residues apart in the sequence; a cluster is defined by a group of two neighbor mutations with N= total reported frequency > 4. Residues showed to display increased activity underlined in col.1.

**Table S2.** RMSD and Overlaps between the Essential Dynamics Modes of 500ns Metatrajectories and the *1NQL* to *3NJP* conformational change.

| STATE (PDB) | Total Time | Variant | Average rMSD | rMSD Max/Min | Overlap (5)<sup>+</sup> | Overlap (10)<sup>+</sup> | Overlap (max)<sup>&</sup> |
| --- | --- | --- | --- | --- | --- | --- | --- |
| **Closed (1NQL)** | 500ns | WT | 8.22 ± 1.74 | 14.00 / 1.20 | 0.81 | 0.85 (0.69) | 0.60 (0.56) |
|  |  | WT+EGF | 6.62 ± 1.75 | 14.25 / 1.06 | 0.86 | 0.91 (0.77) | 0.78 (0.84) |
|  |  | R84K | 8.68 ± 3.02 | 14.93 / 1.34 | 0.77 | 0.91 (0.72) | 0.46 (0.58) |

\* **Overlap (m)** = cumulative overlap ( $\delta_m$ , eq.3) between the first *m* essential modes of the corresponding metatrajectory and the 1NQL to 3NJP transition vector; in brackets: similarity ( $\gamma_m$ , eq.4) between the same modes and ED-ENM Normal Modes. <sup>&</sup> **Overlap (max)** = overlap between the best-aligned *k*-th essential mode ( $\alpha_k$ , eq.3) and the transition vector; in brackets: maximal overlap between an essential mode and an ED-ENM Normal Mode.

**Table S3.** Summary of I-II interdomain hydrogen bonds during 500ns MD meta-trajectories. Note the disappearance of the key Arg84-Cys227 hydrogen bond in R84K. Results are reported as time percentage of the trajectory when bonds are present. Mutated residues in bold; first and second neighbors in italics.

| Domain I-II H-bond |  | WT | R84K | WT+EGF |
| --- | --- | --- | --- | --- |
| ARG198-side | GLU118-side | 95% | 100% | 90% |
| ARG84-side | **CYS227-main** | 45% | 0% | 40% |
| CYS199-main | *THRE187-main* | 31% | 30% | 30% |
| ARG231-side | NGL3-side | 30% | 18% | 30% |
| ALA214-main | GLU118-side | 22% | 31% | 60% |
| ARG198-side | *THRE187-side* | 24% | 23% | 33% |
| *THRE187-side* | *ALA214-main* | 28% | 18% | 38% |
| TYR275-side | GLN8-side | 18% | 11% | 22% |
| ARG198-side | ASP142-main | 20% | 27% | 11% |
| ARG228-side | GLU118-side | 15% | 8% | 0% |

**Table S4.** Summary of II-IV interdomain hydrogen bonds during 500ns MD meta-trajectories. Results are reported as time percentage of the trajectory when bonds are present. Mutated residues in bold; first and second neighbors in italics.

| Domain II-IV H-bond |  | WT | R84K | WT+EGF |
| --- | --- | --- | --- | --- |
| TYR246-main | *MET576-main* | 50% | 10% | 15% |
| TYR246-side | ASP563-side | 13% | 1% | 3% |
| TYR251-side | **ALA573-main** | 5% | 0% | 1% |
| TYR246-side | **GLY574-main** | 5% | 0% | 1% |
| HIS566-main | **TYR251-main** | 3% | 1% | 1% |
| HIS566-side | **THR250-main** | 3% | 2% | 3% |
| *ASN247-side* | **GLU578-side** | 2% | 2% | 4% |
| ASN256-side | **GLU578-side** | 2% | 3% | 1% |
| **GLU578-side/main** | **TYR246-main** | 2% | 1% | 3% |
| ASN256-side | **GLU578/ASN580-side** | 2% | 3% | 1% |
| *ASN579-side* | **THR239-main/side** | 2% | 0% | 0% |
| **ASN580-side** | **THR239-side** | 2% | 0% | 0% |

**Table S5.**
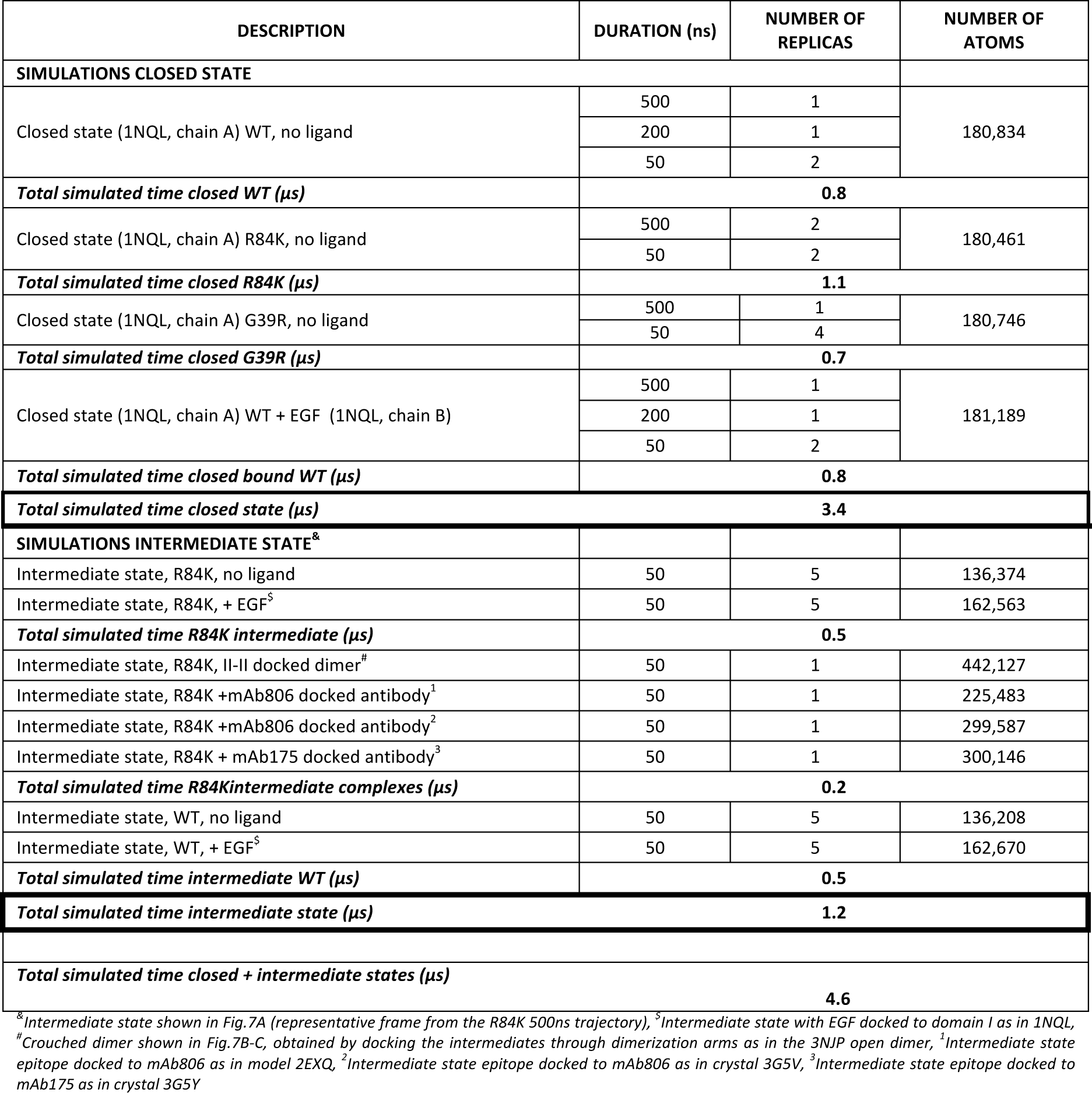
Summary of the MD simulations performed.

